# Proteomics identifies signal peptide features determining the substrate specificity in human Sec62/Sec63-dependent ER protein import

**DOI:** 10.1101/867762

**Authors:** Stefan Schorr, Duy Nguyen, Sarah Haßdenteufel, Nagarjuna Nagaraj, Adolfo Cavalié, Markus Greiner, Petra Weissgerber, Marisa Loi, Adrienne W. Paton, James C. Paton, Maurizio Molinari, Friedrich Förster, Johanna Dudek, Sven Lang, Volkhard Helms, Richard Zimmermann

**Affiliations:** Medical Biochemistry and Molecular Biology, Saarland University, 66421 Homburg, Germany; Center for Bioinformatics, Saarland University, 66041 Saarbrücken, Germany; Max Planck Institute of Biochemistry, Core facility, 82152 Martinsried, Germany; Experimental and Clinical Pharmacology and Toxicology, Saarland University, 66421 Homburg, Germany; Università della Svizzera italiana, Faculty of Biomedical Sciences, Institute for Research in Biomedicine, CH-6500 Bellinzona, Switzerland; Research Centre for Infectious Diseases, University of Adelaide, South Australia 5005, Australia; Bijvoet Center for Biomolecular Research, Utrecht University, 3584 CH Utrecht, The Netherlands; Weizman Institute of Science, Rehovot 7610001, Israel

**Author notes:** Corresponding authors Sven Lang, Medical Biochemistry and Molecular Biology, Saarland University, 66421 Homburg, Germany, Richard Zimmermann, Medical Biochemistry and Molecular Biology, Saarland University, 66421 Homburg, Germany. These authors contributed equally to this work (Stefan Schorr, Duy Nguyen). **Abbreviations**: BiP, immunoglobulin heavy-chain binding protein; C-region, carboxy-terminal region of SP; ER, endoplasmic reticulum; GO, gene ontology; H-region, hydrophobic region of SP; N-region, amino-terminal region of SP; PLD, Polycystic Liver Disease; SP, signal peptide; SRP, signal recognition particle; TMH, SP-equivalent transmembrane helix; TRAP, translocon-associated protein; UTR, untranslated region; WT, wild type.

**Keywords:** Endoplasmic reticulum, protein import, Sec61-channel, Sec62, Sec63

## Abstract

In mammalian cells one-third of all polypeptides are integrated into the membrane or translocated into the lumen of the endoplasmic reticulum (ER) via the Sec61-channel. While the Sec61-complex facilitates ER-import of most precursor polypeptides, the Sec61-associated Sec62/Sec63-complex supports ER-import in a substrate-specific manner. So far, mainly posttranslationally imported precursors and the two cotranslationally imported precursors of ERj3 and prion protein were found to depend on the Sec62/Sec63-complex *in vitro*. Therefore, we determined the rules for engagement of Sec62/Sec63 in ER-import in intact human cells using a recently established unbiased proteomics approach. In addition to confirming ERj3, we identified twenty-two novel Sec62/Sec63-substrates under these *in vivo*-like conditions. As a common feature, those previously unknown substrates share signal peptides with comparatively longer but less hydrophobic H-region and lower C-region polarity. Further analyses with four substrates, and ERj3 in particular, revealed the combination of a slowly-gating signal peptide and a downstream translocation-disruptive positively charged cluster of amino acid residues as decisive for the Sec62-/Sec63-requirement. In the case of ERj3, these features were found to be responsible for an additional BiP-requirement and to correlate with sensitivity towards the Sec61-channel inhibitor CAM741. Thus, the human Sec62/Sec63-complex may support Sec61-channel opening for precursor polypeptides with slowly-gating signal peptides by direct interaction with the cytosolic amino-terminal peptide of Sec61α or via recruitment of BiP and its interaction with the ER-lumenal loop 7 of Sec61α. These novel insights into the mechanism of human ER protein import contribute to our understanding of the etiology of *SEC63*-linked Polycystic Liver Disease.

**Databases:** The mass spectrometry proteomics data have been deposited to the ProteomeXchange Consortium via the PRIDE partner repository (http://www.ebi.ac.uk/pride/archive/projects/Identifiers) with the dataset identifiers: PXD008178, PXD011993, and PXD012078. Supplementary information was deposited at Mendeley Data under the DOI:10.17632/6s5hn73jcv.1 (http://dx.doi.or/10.17632/6s5hn73jcv.1).

## Introduction

The endoplasmic reticulum (ER) of mammalian cells handles most secreted as well as soluble and membrane proteins of the secretory pathway [1–5]. Typically, protein translocation across or insertion into the ER-membrane is linked to protein synthesis (cotranslational) and mediated by i) signal peptides (SP) or SP-equivalent transmembrane helices (TMH) in the precursors [5–7], as well as ii) a targeting system, which involves the cytosolic signal recognition particle (SRP) plus the SRP-receptor in the ER-membrane [8–10], and iii) a polypeptide-conducting channel in the ER-membrane, the heterotrimeric Sec61-complex [11]. Elegant *in vitro* experiments demonstrated that at least for some precursor polypeptides (such as bovine preprolactin) these components are sufficient for ER-import [12]. In contrast, other precursor polypeptides with SP or TMH additionally rely on Sec61-auxiliary and -associated membrane components, such as the heterotetrameric translocon-associated protein complex (TRAP) [13, 14] or the heterodimeric Sec62/Sec63-complex [15–18]. Both complexes support Sec61-channel opening in the case of precursors with inefficiently-gating SP plus, in some cases, detrimental features in their mature regions [14, 17–22]. A proteomic identification of TRAP-dependent precursors characterized a high glycine- and proline-content and/or low hydrophobicity of the SP as distinguishing features for TRAP-dependence [21]. In combination with structural analysis [22], these results suggested a scenario, where TRAPγ recognizes TRAP-clients on the cytosolic ER-surface and Sec61-channel opening is mediated by direct interaction of the ER-lumenal domains of the TRAP α- and β-subunits with the ER-lumenal loop 5 of Sec61α [21]. The Sec62/Sec63-complex can support Sec61-channel opening either by an yet-undefined mechanism on its own [19] or in cooperation with the ER-lumenal BiP, which can be recruited to the Sec61-complex via Sec63 and directly interact with the ER-lumenal loop 7 of Sec61α [19, 23]. So far, the substrate spectrum of the mammalian Sec62/Sec63-complex as well as its rules of engagement for substrate recognition and Sec61-channel opening remained ill-defined. This gap was not closed by the recent structural analysis of the yeast heptameric SEC-complex, which includes aside from the trimeric Sec61-complex the tetrameric Sec62/Sec63/Sec71/Sec72-complex [24, 25]. Of note, the yeast SEC-complex is supposedly involved only in posttranslational and SRP-independent protein import into the ER and the additional components, Sec71 and Sec72, are without mammalian orthologs.

SP for ER protein import, typically, comprise around 25 amino acid residues and have a tripartite structure with a positively charged N-region, a central H-region, and a slightly polar C-region [6, 26]. They target presecretory proteins to the Sec61-complex and facilitate the opening of an aqueous channel within the Sec61-complex for passage of the polypeptide to the ER-lumen [12, 27–30]. TMH are similar to SP with respect to structure and function, except for the positioning of positively charged amino acid residues, which can be up- or downstream of the central hydrophobic region and determine its insertion/orientation in the ER membrane [6, 7, 26–30]. *A priori*, SP and TMH can insert into the Sec61-channel in a head-on (N_ER-lumen_-C_cytosol_) or a loop (N_cytosol_-C_ER-lumen_) configuration.

According to *in vitro* experiments with insect (preprocecropin A) and human model proteins (preproapelin, prestatherin), the import of presecretory proteins with short and apolar SP and an over-all length of below 100 amino acid residues (small precursor proteins) into the mammalian endoplasmic reticulum (ER) occurs posttranslationally [17-19, 31-33], and involves all known targeting mechanisms [19] and Sec62 plus Sec63 for Sec61-channel opening [17–19, 33]. In the case of preprocecropin A, posttranslational ER-import has also been shown in human cells [32]. Furthermore, Sec62/Sec63-dependence of small human presecretory proteins was confirmed in human cells [17]. However, human Sec62 and Sec63 can also be involved in cotranslational import of precursor proteins with a size of more than 100 amino acid residues into the mammalian ER (i.e. the precursors of the ER-lumenal protein ERj3 with 358 amino acid residues, and the GPI-anchored prion protein with 253 amino acid residues), either in the standard *in vitro* import assay [18, 20] or with corresponding nascent polypeptide chains [29]. Under the latter conditions, it was shown that these two precursors are slow in inducing Sec61-channel opening and, maybe therefore, recruit Sec62 and Sec63 relatively late in their synthesis to the channel [29]. In the case of the prion protein precursor, Sec62/Sec63-dependent ER-import has also been demonstrated in human cells [34]. In addition, it was observed in the *in vitro* import assays that Sec63-dependence of preproapelin and the precursors of ERj3 and prion protein is related to gating of the Sec61-channel to the open state and correlates with BiP’s action, which was linked to the combination of the respective SP plus, in case of preproapelin (_37_RRK) and the prion protein precursor (_1_KKRPK), a positively charged cluster in the mature region [19, 20, 23]. Furthermore, loss of Sec63 protein function in the liver of a subset of human patients, who suffer from Autosomal Dominant Polycystic Liver Disease (PLD) (OMIM174050), was also interpreted in light of a substrate specific function of Sec63 in ER protein import [35]. Interestingly, Sec63-dependence of ERj3 import into the ER was indirectly confirmed in murine *SEC63* null cells, which were generated as an animal model for the human disease [35]. These *SEC63^-/-^* cells lacked ERj3 while the levels of various other ER proteins were unchanged compared to murine *SEC63^+/+^* cells [18].

Here we address the questions of which precursor polypeptides depend on Sec62 and Sec63 in their ER-import in intact human cells under steady-state conditions and what common features these precursor polypeptides share. A recently established unbiased proteomics approach [21] identified a total of 36 precursors as Sec62/Sec63-clients, 23 of which share SP with longer but less hydrophobic H-region and lower C-region polarity than average. Independent validation experiments characterized a combination of a slowly gating SP and the detrimental effect of a positively charged cluster downstream of the SP as decisive for the Sec62-/Sec63-plus BiP-requirement of ERj3 in human cells.

## Results

### Substrate specificity of Sec62/Sec63 in ER protein import in HeLa cells

We first addressed the substrate spectrum of human Sec62 and Sec63 by the established combination of siRNA-mediated depletion of Sec62 or Sec63 in HeLa cells, label-free quantitative proteomics, and differential protein abundance analysis [21]. The cells were treated with two sets of different siRNAs that target either *SEC62* or *SEC63* or with a non-targeting (control) siRNA in triplicates for 96 h. It was previously established that this gene silencing method resulted in >90% Sec62- or Sec63-depletion, without significantly affecting cell growth, cell viability, or cell/ER morphology [18, 19].

After Sec62 depletion in two independent experiments, up to 6,686 different proteins were quantitatively characterized by MS, 4,819 of which were detected in all samples (Fig. 1, Table 1, Table S1-5, http://www.ebi.ac.uk/pride/archive/projects/Identifiers PXD008178, PXD012078). Applying the established statistical analysis [21], we found that transient Sec62 depletion significantly affected the steady-state levels of 351 proteins: 155 negatively and 196 positively (q<0.05) (Table S2, 3). As expected [18], Sec62 itself was negatively affected (Fig. 1A), which was confirmed by Western blot (Fig. 1B). Similar to the total quantified proteome, ∼25% of the negatively affected proteins were assigned to organelles of the endocytic and exocytic pathway by GO term analysis (Fig. 1C, large pies). We also did not detect significant enrichment of proteins with SP, N-glycosylated proteins, and membrane proteins (Fig. 1C, small pies). Nevertheless, the identified precursors included 18 proteins with cleavable SP and six proteins with TMH (Table S4, 5). ERj3, product of the *DNAJB11* gene, was negatively affected (Fig. 1A), which confirms the expectation that non-transported precursors of Sec62-clients are degraded by the proteasome upon its depletion.

**Fig. 1.**
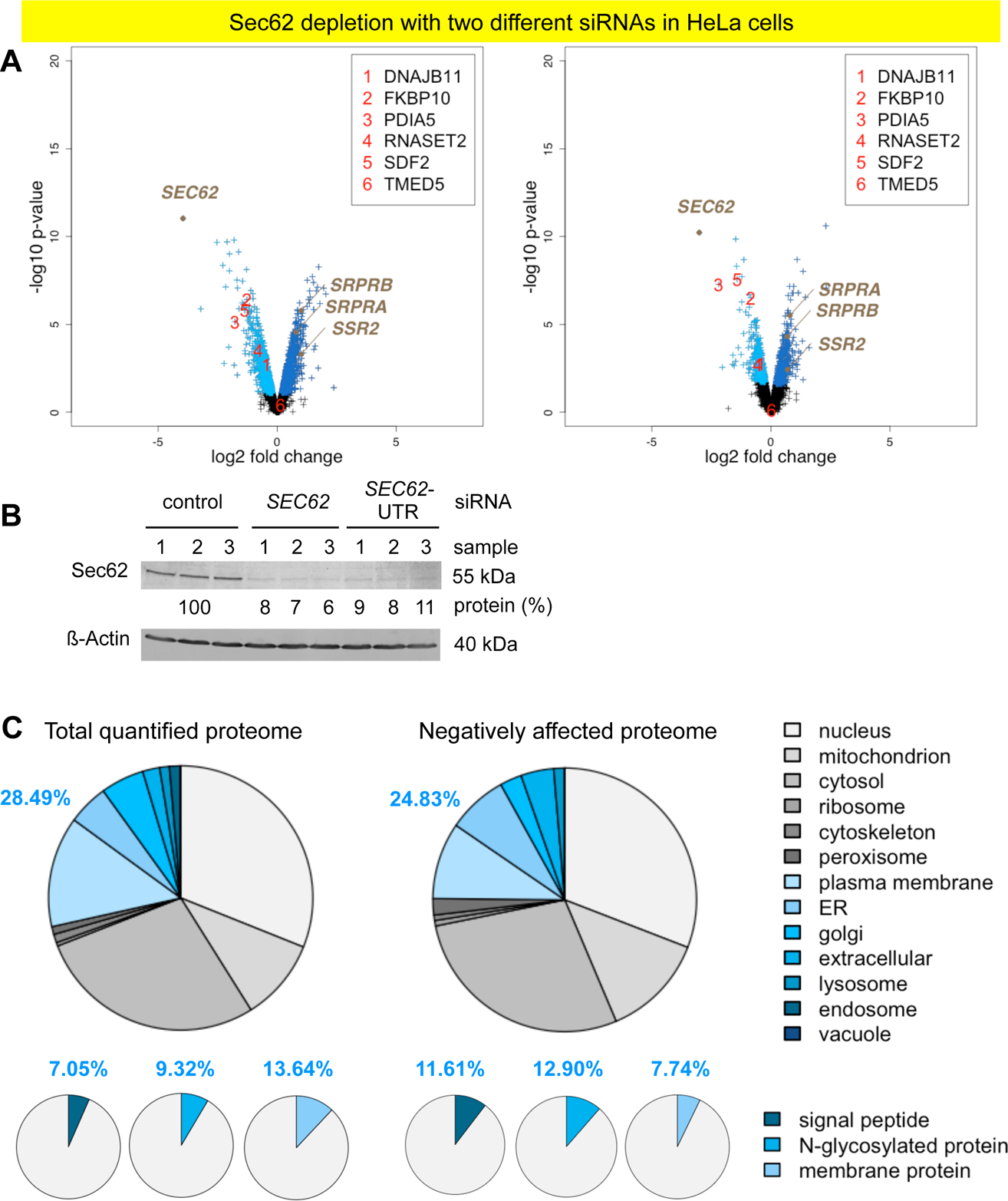
Identification of Sec62-clients and compensatory mechanisms by knock-down of Sec62 in HeLa cells. The experimental strategy was as follows: two sets of three replicates from two independent experiments for each cell type; label-free quantitative proteomic analysis; and differential protein abundance analysis to identify negatively affected proteins (i.e. clients) and positively affected proteins (i.e. compensatory mechanisms). (A) Differentially affected proteins were characterized by the mean difference of their intensities plotted against the respective *p* values in volcano plots. Six negatively affected proteins, representing the overlap between negatively affected SP-containing proteins after Sec62-depletion in HeLa and *SEC62* knock-out in HEK293 cells plus the TRAP-client TMED5 (Table 2), are highlighted with red numbers. (B) Knock-down efficiencies were evaluated by Western blot. Results are presented as % of residual protein levels (normalized to ß-actin) relative to control, which was set to 100%. (C) Protein annotations of SP, membrane location, and N-glycosylation in humans were extracted from UniProtKB and used to determine the enrichment of Gene Ontology annotations among the secondarily affected proteins. Notably, there was no indication for activation of the unfolded protein response (UPR) in the course of the 96 h knock-down, i.e. related terms did not come up as enriched GO terms in the analysis of the positively affected proteins and typical UPR-regulated genes such as *HSPA5* (coding for BiP) and *HYOU1* (coding for Grp170) were not up-regulated (Table S3).

**Table 1.** Statistics of the identification of Sec62- and Sec63-clients in comparison to Sec61α1- and TRAP-clients, respectively.

| Proteins | SEC61A1 | TRAP | SEC62 | SEC63 | SEC62HEK | SEC63HEK |
| --- | --- | --- | --- | --- | --- | --- |
| Quantified proteins | 7212 | 7670 | 6686 | 6655 | 6195 | 6195 |
| Statistically analyzed proteins | 5129 | 5911 | 4819 | 6655 | 6195 | 6195 |
| representing the secretory pathway (%) | 26 | 27 | 28 | 28 | 26 | 26 |
| Proteins with SP (%) | 6 | 7 | 7 | 7 | 5 | 5 |
| N-Glycoproteins (%) | 8 | 8 | 9 | 9 | 7 | 7 |
| Membrane proteins (%) | 12 | 13 | 14 | 14 | 12 | 12 |
| Positively affected proteins | 342 | 77 | 196 | 13 | 121 | 103 |
| Negatively affected proteins | 482 | 180 | 155 | 21 | 208 | 199 |
| representing the secretory pathway (%) | 61 | 40 | 25 | 50 | 48 | 37 |
| Negatively affected proteins with SP (%) | 41 | 22 | 12 | 14 | 30 | 11 |
| Negatively affected N-glycoproteins (%) | 45 | 23 | 13 | 24 | 33 | 18 |
| Negatively affected membrane proteins (%) | 36 | 26 | 8 | 38 | 21 | 21 |
| Negatively affected proteins with SP | 197 | 38 | 18 | 3 | 62 | 21 |
| including N-glycoproteins | 158 | 28 | 12 | 2 | 50 | 16 |
| corresponding to % | 80 | 74 | 67 | 67 | 81 | 76 |
| including membrane proteins | 77 | 19 | 2 | 2 | 18 | 9 |
| corresponding to % | 39 | 50 | 11 | 67 | 29 | 43 |
| Negatively affected proteins with TMH | 98 | 22 | 6 | 6 | 22 | 29 |
| including N-glycoproteins | 56 | 11 | 5 | 3 | 8 | 14 |
| corresponding to % | 57 | 50 | 83 | 50 | 36 | 48 |

Comparatively low numbers are highlighted in yellow.

The proteins positively affected by transient Sec62 depletion included both SRP-receptor subunits (SRPRA, SRPRB) and various ubiquitin-conjugating enzymes (such as MID1). This is consistent with the temporary, cytosolic accumulation of precursors before proteasomal degradation and reminiscent of the previously found compensatory mechanisms after Sec61- and TRAP-complex depletion (Table S3) [21]. Interestingly, the TRAP-complex ß-subunit (coded by the *SSR2* gene) was also positively affected by transient Sec62 depletion, which may suggest overlapping functions of Sec62 and TRAP in targeting or Sec61-channel gating. These short-term compensatory mechanisms may have contributed to the comparatively low number of negatively affected proteins.

After analogous Sec63-depletion, 6,655 different proteins were quantitatively characterized by MS, which were detected in all samples (Fig. 2, Table 1, Table S6-9). Here, we found that Sec63-depletion significantly affected the steady-state levels of only 34 proteins: 21 negatively and 13 positively (q<0.05) (Table S7, 8). As expected Sec63 itself was negatively affected (Fig. 2A) and confirmed (Fig. 2B). GO terms assigned 50% of the negatively affected proteins to organelles of the endocytic and exocytic pathways (Fig. 2C, large pies). We also detected significant enrichment of proteins with SP (2-fold), N-glycosylated proteins (2.6-fold), and membrane proteins (2.8-fold) (Fig. 2C, small pies). However, the identified precursors included only four proteins with cleavable SP and six proteins with TMH and only one of the proteins with SP had also been negatively affected by Sec62-depletion, TGFBI (Table S9, Table 2). Upon closer inspection of the potential overlap between *SEC62* and *SEC63* silencing in HeLa cells, four additional precursor polypeptides with SP were negatively affected by Sec63-depletion in HeLa cells, which did not meet the stringent significance threshold (ERj3, MAGT1, PDIA5, SDF2) (Table 2). Thus, at least these five precursors depend on the Sec62/Sec63-complex in HeLa cells for their ER import.

**Fig. 2.**
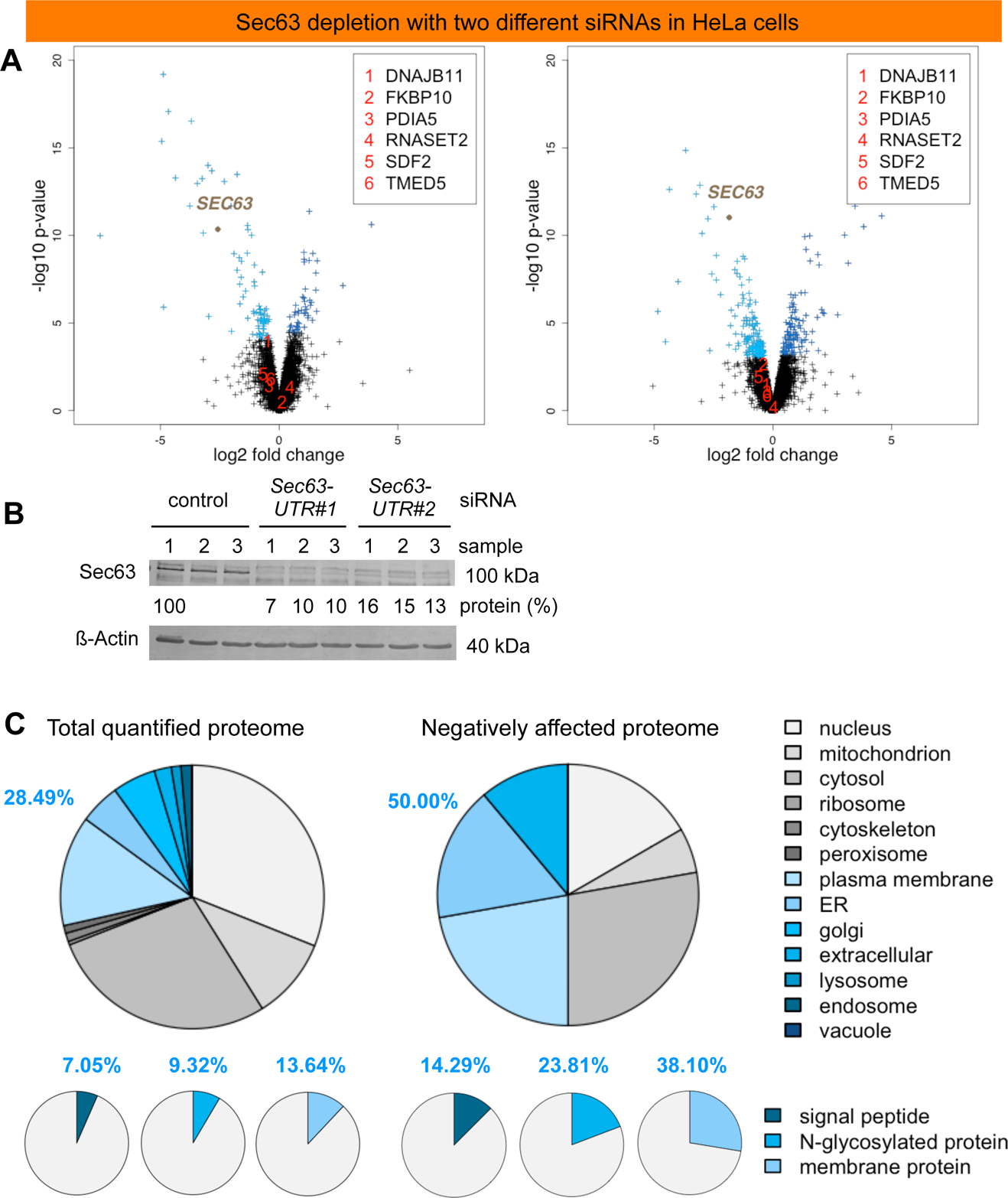
Identification of Sec63-clients and compensatory mechanisms by knock-down of Sec63 in HeLa cells. The experimental strategy was as follows: three replicates for each cell type; label-free quantitative proteomic analysis; and differential protein abundance analysis to identify negatively affected proteins (i.e. clients) and positively affected proteins (i.e. compensatory mechanisms). (A) Differentially affected proteins were characterized by the mean difference of their intensities plotted against the respective *p* values in volcano plots. Six negatively affected proteins of Sec62 depletion are indicated with red numbers. (B) Knock-down efficiencies were evaluated by Western blot. Results are presented as % of residual protein levels (normalized to ß-actin) relative to control, which was set to 100%. (C) Protein annotations of SP, membrane location, and N-glycosylation in humans were extracted from UniProtKB and used to determine the enrichment of Gene Ontology annotations among the secondarily affected proteins. We note that the positively affected proteins included a signal peptidase complex subunit, but no ubiquitin-conjugating enzymes, consistent with the fact that there were only a couple of proteins negatively affected and, therefore, no significant accumulation of precursor polypeptides in the cytosol after partial Sec63 depletion for 96 h (Table S8).

**Table 2.**
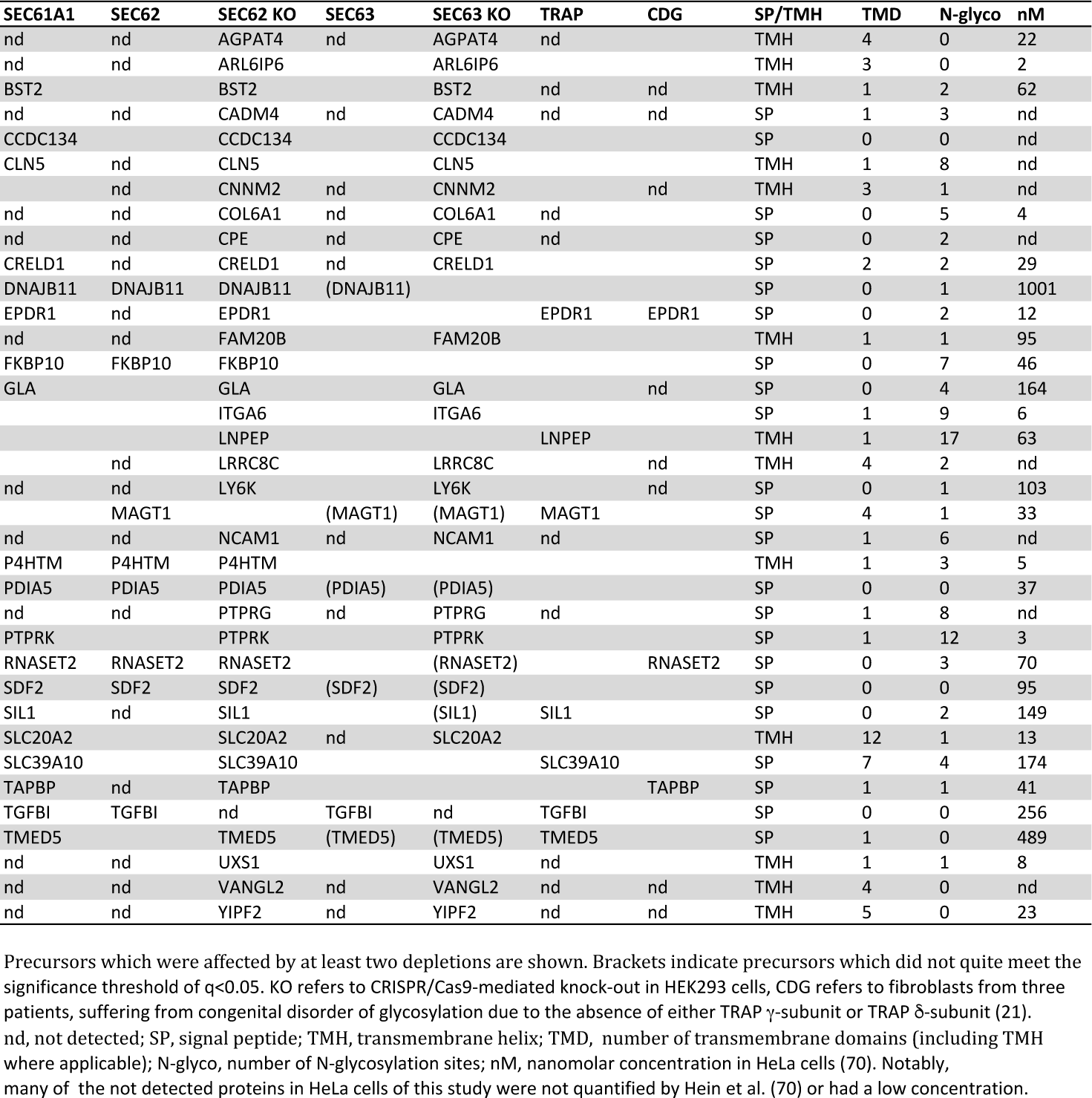
Overlap of negatively affected precursor polypeptides after transient or permanent depletion of Sec61α, Sec62, Sec63, or TRAP from human cells.

### Substrate specificity of Sec62/Sec63 in ER protein import in HEK293 cells

To possibly identify additional substrates by permanent depletion, we performed similar analyses after Sec62- or Sec63-depletion, using the respective CRISPR/Cas9 treated HEK293 cells compared to HEK293 control cells [36].

After Sec62 depletion in HEK293 cells, 6195 different proteins were quantitatively characterized by MS, which were detected in all samples (Fig. 3, Table S10-14, http://www.ebi.ac.uk/pride/archive/projects/Identifier PXD011993). Applying the statistical analysis, we found that Sec62 depletion significantly affected the steady-state levels of 329 proteins: 208 negatively and 121 positively (q<0.05) (Table S11, 12). Sec62 itself was negatively affected (Fig. 3A), which was confirmed by Western blot (Fig. 3B). Of the negatively affected proteins, GO terms assigned ∼48% to organelles of the endocytic and exocytic pathways (Fig. 3C). We also detected significant enrichment of proteins with SP (5.5-fold), N-glycosylated proteins (4.5-fold), and membrane proteins (1.8-fold) (Fig. 3C). The identified precursors included 62 proteins with cleavable SP and 22 proteins with TMH. As expected, ERj3 was negatively affected (Table S13, 14). Notably, six precursor polypeptides were negatively affected by transient as well as permanent Sec62-depletion, including five precursors with SP (ERj3, FKBP10, PDIA5, RNaseT2, SDF2) and one protein with TMH (P4HTM) (Fig. 3D, Table 2). The positively affected proteins included a cytosolic molecular chaperone (HSPB1) and several cytosolic ubiquitin-conjugating enzymes (RNF31, UBE4B, WWP2), (Table S12), consistent with cytosolic accumulation of precursors.

**Fig. 3.**
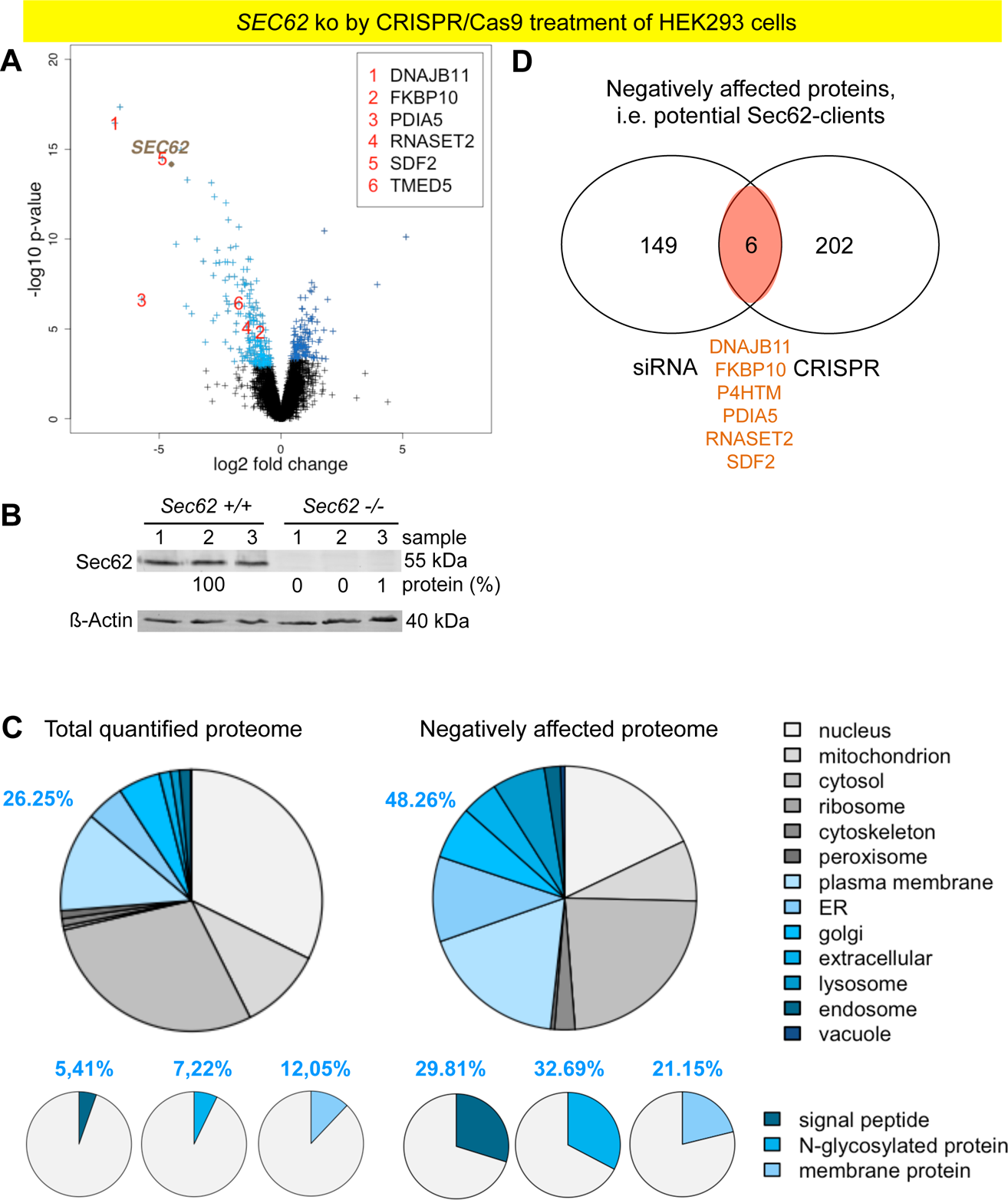
Identification of Sec62-clients and compensatory proteins by CRISPR/Cas9-mediated Sec62-depletion in HEK293 cells. (A) The experimental strategy was as described in Fig. 2. Six negatively affected proteins of Sec62 depletion in HeLa cells are indicated with red numbers. (B) Knock-out efficiencies were evaluated by Western blot. Results are presented as % of residual protein levels (normalized to ß-actin) relative to control, which was set to 100%. (C) Protein annotations of SP, membrane location, and N-glycosylation in humans were extracted from UniProtKB, and used to determine the enrichment of Gene Ontology annotations among the secondarily affected proteins. (D) Venn diagram for the overlap between negatively affected proteins after Sec62-depletion in HeLa and *SEC62* knock-out in HEK293 cells. Notably, there was a slight UPR activation, resulting in overproduction of ER-lumenal chaperones (BiP, Grp94) and components involved in ER associated protein degradation (EDEM3, SPPL2A), which we attribute to the almost complete absence of the BiP co-chaperone ERj3 (Table S12).

After Sec63 depletion in HEK293 cells, the above-mentioned 6195 different proteins were quantitatively characterized by MS, which were detected in all samples (Fig. 4, Table S15-19). Here, we found that Sec63-depletion significantly affected the steady-state levels of 302 proteins: 199 negatively and 103 positively (Table S16, 17). Sec63 itself was negatively affected (Fig. 4A), which was supported by Western blot (Fig. 4B). GO terms assigned ∼37% of the negatively affected proteins to organelles of the endocytic and exocytic pathways (Fig. 4C). We also detected significant enrichment of proteins with SP (1.9-fold), N-glycosylated proteins (2.4-fold), and membrane proteins (1.8-fold) (Fig. 4C). The identified precursors included 21 proteins with cleavable SP and 29 proteins with TMH (Table S18, 19). Notably, 22 precursor polypeptides were negatively affected by Sec62 as well as Sec63 knock-down, 11 each with SP and TMH (Fig. 4D, Table 2). ERj3 was not negatively affected under these conditions. However, the Sec63-dependence of ERj3 import into the ER in intact cells had previously been observed for murine *SEC63* null cells [18, 35]. The positively affected proteins included cytosolic molecular chaperones (HSPA2, HSPB1) and a single ubiquitin-conjugating enzyme (Listerin).

**Fig. 4.**
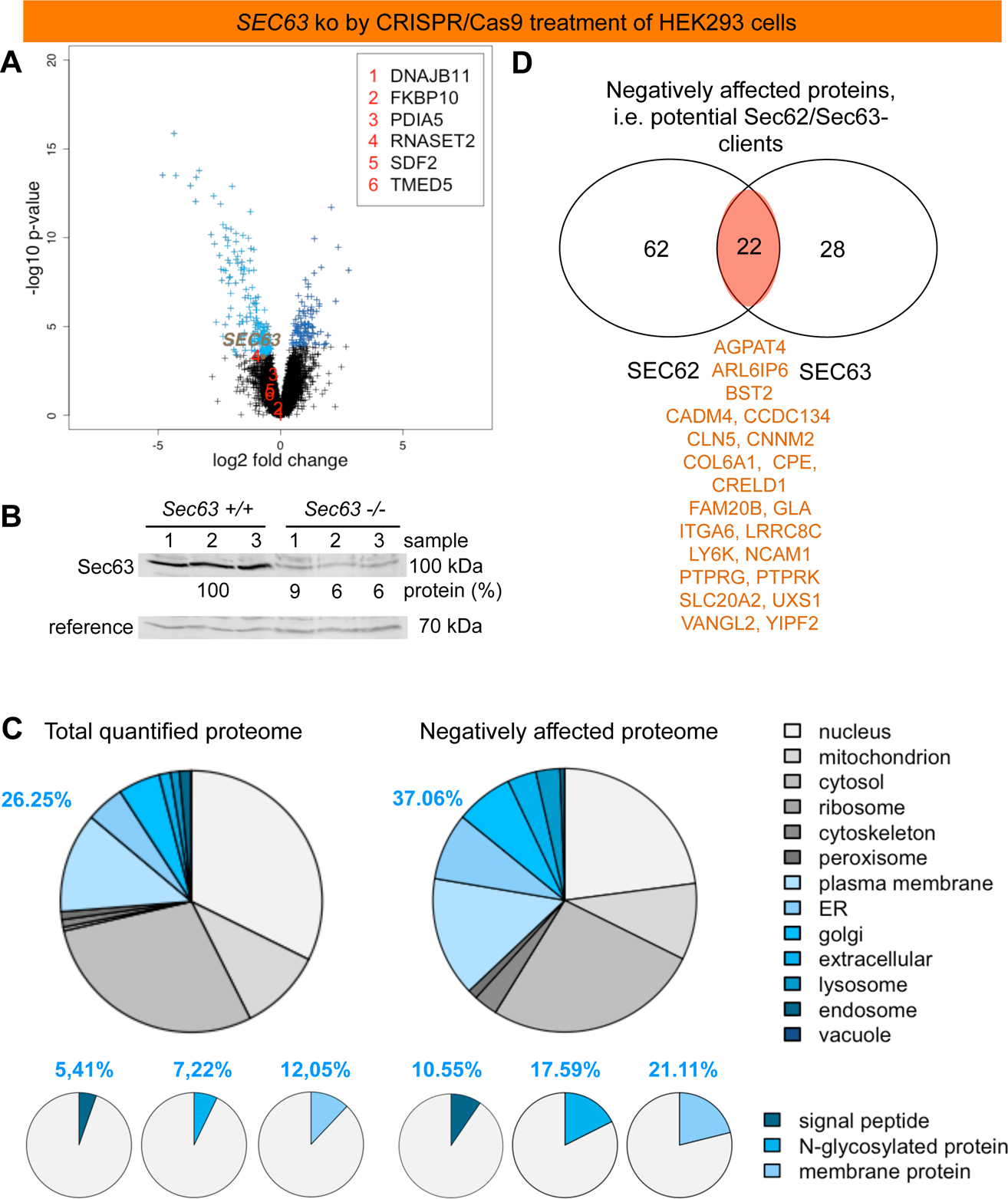
Identification of Sec63-clients and compensatory proteins by CRISPR/Cas9-mediated Sec63-depletion in HEK293 cells. (A) The experimental strategy was as described in Fig. 2. Six negatively affected proteins of Sec62 depletion in HeLa and HEK293 cells are indicated with red numbers. (B) Knock-out efficiencies were evaluated by Western blot. Results are presented as % of residual protein levels (normalized to an unknown cross-reacting protein, termed reference) relative to control, which was set to 100%. (C) Protein annotations of SP, membrane location, and N-glycosylation in humans were extracted from UniProtKB, and used to determine the enrichment of Gene Ontology annotations among the secondarily affected proteins. (D) Venn diagram for the overlap between negatively affected proteins after Sec62- and Sec63-depletion in HEK293 cells. We note that there was no indication for UPR activation (Table S17),

### Validation of Sec6/Se63 substrates

To validate the proteomic data on Sec62/Sec63 substrates, we conducted independent silencing and Western blot experiments with the *SEC62*- and the *SEC63*-UTR-targeting siRNA in HeLa cells for four SP-containing candidates (ERj3, FKBP10, PDIA5, RNaseT2), representing N-glycosylated as well as non-glycosylated proteins and proteins with different abundances in HeLa cells (Table 2). Similar to the proteomic experiments, the cells were treated with targeting or non-targeting siRNA for 96h and analyzed by Western blot. Western blot analysis confirmed ERj3, FKBP10, PDIA5, plus RNaseT2 as Sec62-dependent, and ERj3, PDIA5, plus RNaseT2 as Sec63-dependent (Fig. 5A). FKBP10 was not affected by Sec63 depletion. Thus, the Western blots fully confirmed the proteomic analysis (Table 2).

**Fig. 5.**
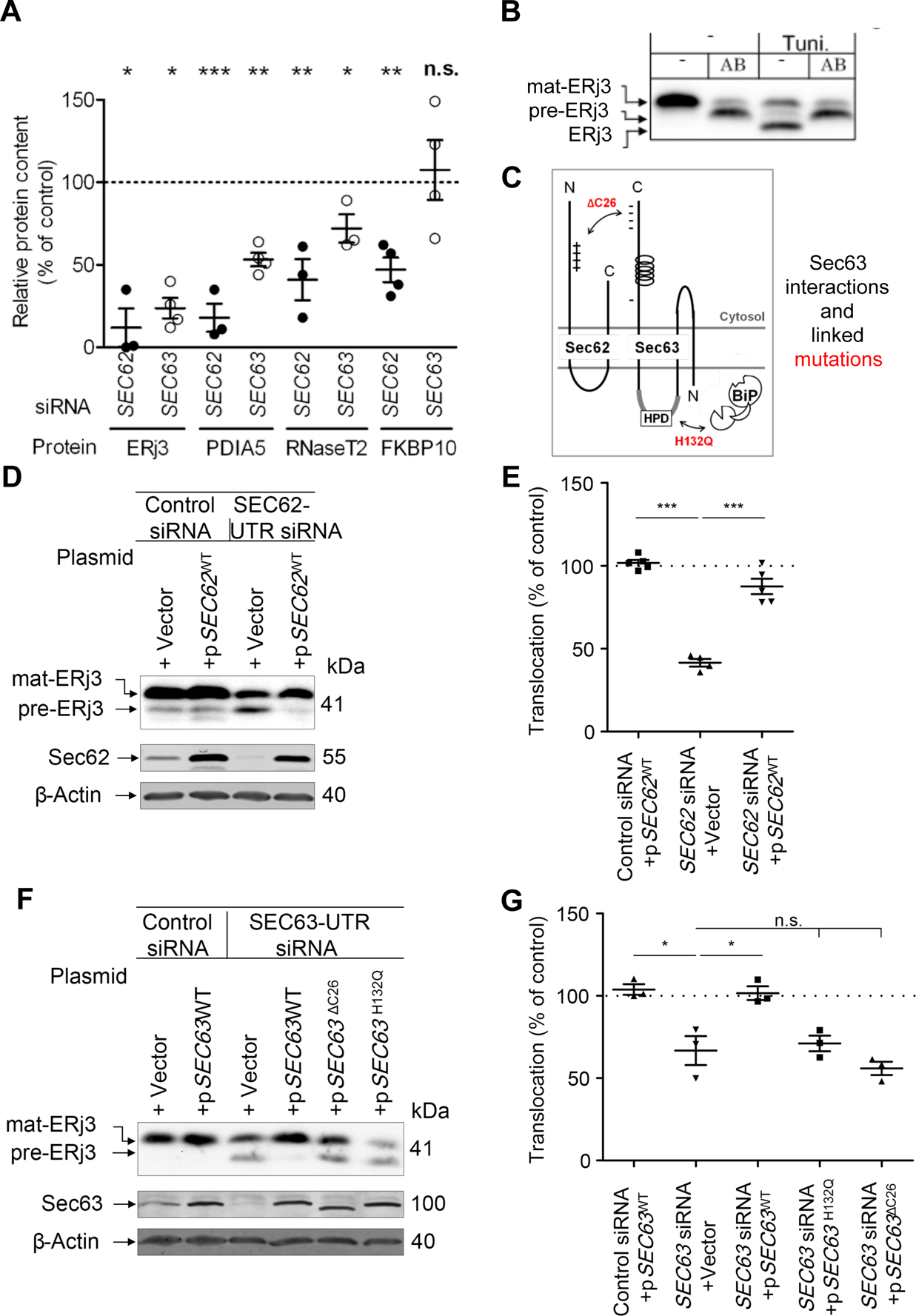
Validation of Sec62- and Sec63-clients by Western blot. (A) To validate the proteomic analysis for four Sec62- and Sec63-clients, HeLa cells were treated with *SEC62*- or *SEC63-* UTR-targeting siRNA or control siRNA and subjected to SDS-PAGE and Western blot. The dot plots depict steady state protein levels, calculated as relative protein content in the targeting siRNA sample with the individual control siRNA sample set to 100% (normalized to ß-actin). Statistical analysis and graphical representation are described in Experimental Procedures. (B) For further validation, one client was analyzed under conditions of its short-term overexpression. To visualize all potential ERj3 forms, human *ERJ3* was expressed in HeLa cells, which had been treated with a transport inhibitor (AB, see below), in the presence of MG132 (10 µM) and in the presence or absence of Tunicamycin (Tuni., 2 µg/ml). The cells were analyzed by SDS-PAGE and Western blot. The relevant section from a representative blot is shown. (C) The cartoon depicts relevant Sec63-interactions and Sec63 mutants (in red) used in the complementation assay. (D-G) Human *ERJ3* was over-expressed in HeLa cells, which had been treated with *SEC62*-UTR-targeting siRNA (D, E), or *SEC63-*UTR-targeting siRNA (F, G), or control siRNA, plus control vector, or *SEC62*- or *SEC63*-expression plasmid, in the presence of MG132. The consequences of depletion and plasmid complementation were analyzed by SDS-PAGE and Western blot for the respective target protein and the model protein ERj3, ß-actin served as a control. Representative blots are shown in D and F. (E, G) Dot plots depict relative pre-ERj3 translocation efficiencies, calculated as the proportion of N-glycosylation of the total amount of synthesized pre-ERj3 with the individual control sample set to 100%. Statistical analysis and graphical representation are described in Experimental Procedures.

For further validation of the proteomic analysis, the effect of Sec62- and Sec63-depletion on ER-import of pre-ERj3 was analyzed in intact HeLa cells under established conditions of pre-ERj3 short-term overproduction and simultaneous proteasome inhibition [21]. HeLa cells were treated with non-targeting siRNA or *SEC62*-UTR- or *SEC63*-UTR-targeting siRNA for 72 h, incubated in the presence of the pre-ERj3 expression plasmid for additional 24 h, and subjected to SDS-PAGE and Western blot. During the last 8 h of the incubation, the proteasome inhibitor MG132 was present. Notably, the SDS-PAGE was able to separate precursor (pre-ERj3), from precursor that was processed by signal peptidase (ERj3), and from the mature i.e. processed plus N-glycosylated protein (mat-ERj3) (Fig. 5B). According to Western blots, Sec62-and Sec63-depletion caused accumulation of pre-ERj3 and concomitant inhibition of ERj3 import into the ER, measured as absence of ERj3 as well as mat-ERj3 (Fig. 5C-G). As a control for the specificity of the siRNA effects, complementation of the siRNA knock-down by expression of the *SEC62* or *SEC63* cDNA was used. In these rescue experiments, *SEC62* or *SEC63* expression plasmid or the vector control were transfected 48 h after the first siRNA treatment. Expression of siRNA-resistant *SEC62* or *SEC63* prevented accumulation of pre-ERj3 and concomitant inhibition of ERj3 import into the ER, i.e. rescued the phenotype of Sec62- or Sec63-depletion (Fig. 5D-G). Thus, pre-ERj3 is a *bona fide* Sec62-and Sec63-client.

The successful complementation of *SEC63*-UTR siRNA phenotypes by *SEC63* cDNA expression allowed the analysis of Sec63 mutant variants with a C-terminal truncation of 26 several negatively charged amino acids in the cytosolic domain (∆C26), which prevents Sec62-interaction, or with a point mutation (H132Q) in the characteristic HPD motif in the ER-lumenal J-domain, which suppresses productive BiP-interaction (Fig. 5C) [19, 37]. When the effects of Sec63H132Q or Sec63ΔC26 overproduction were analyzed in the presence of *SEC63*-UTR siRNA, both mutant variants failed to rescue the phenotype of Sec63-depletion (Fig. 5F, G). The failure of the mutant with a deletion of 26 negatively charged amino acids indirectly confirmed the observed role of Sec62 in pre-ERj3 ER-import and suggested that Sec62 and Sec63 are acting together in a complex in the ER-import of pre-ERj3. The failed rescue of the H132Q mutant suggested that BiP is involved in the import of pre-ERj3 into the ER of human cells as well (see next section).

Next, we asked where in the cell pre-ERj3 is accumulating in the absence of Sec62 and Sec63. First, pre-ERj3 accumulating HeLa cells, which were depleted of Sec62, were converted to semi-permeabilized cells by treatment with digitonin and then incubated in the absence or presence of proteases. Accumulated pre-ERj3 was protease-sensitive in the absence of Triton X-100, while mat-ERj3 was protease-resistant in the absence of Triton X-100 and protease-sensitive in its presence (Fig. 6A, B). Thus, mature ERj3 had reached the ER-lumen in the presence of Sec62, while in the absence of Sec62 accumulating pre-ERj3 remained in the cytosol, at least partially. In the second approach, the semi-permeabilized cells were subjected to extraction at pH 11.5, which is employed to differentiate between soluble and phospholipid bilayer-integrated proteins. Mature ERj3 was solubilized at pH 11.5, while pre-ERj3 was resistant towards alkaline extraction (Fig. 6C, D). Thus pre-ERj3, which accumulated in the absence of Sec62, may have been integrated into the ER membrane via its un-cleaved SP, which would be consistent with the fact that no ERj3 (i.e. precursor that was processed by signal peptidase but not N-glycosylated) was observed under these conditions. This may indicate that in Sec62 absence, the SP of pre-ERj3 either cannot insert into the Sec61-channel in the productive loop configuration or it inserts head-on and cannot make the flip turn. Sec63-depletion phenocopied this result (Fig. 6C, D).

**Fig. 6.**
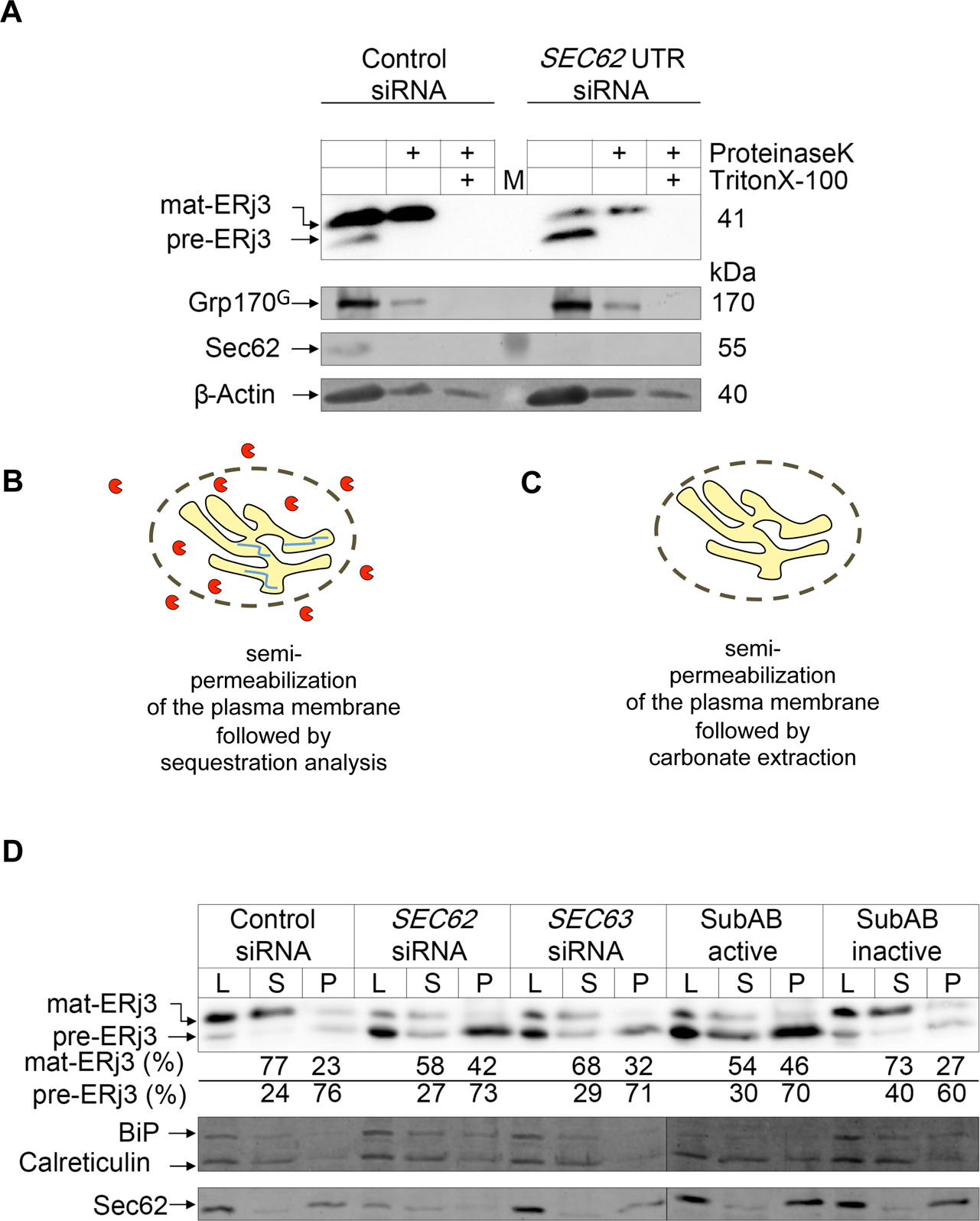
Cellular localization of pre-ERj3 in the absence of Sec62, Sec63, or BiP. (A-D) Human *ERJ3* was expressed in HeLa cells, which had been treated with *SEC62*-UTR-targeting siRNA (A, D), or *SEC63-*UTR-targeting siRNA (D), or control siRNA (A, D), or with active or inactive SubAB, in the presence of MG132 (D). (A, D) After semi-permeabilization of the cells, the consequences of depletion were analyzed by either protease accessibility- (A, B) or alkaline extraction- (C, D) analysis and followed by SDS-PAGE and Western blot for the respective target protein, the model protein ERj3, and Calreticulin and the N-glycosylated Grp170^G^ as ER lumenal marker proteins, ß-actin served as a control. In protease-accessibility the proteases trypsin and proteinase K were each present at final concentrations of 50 µg/ml. (B, C) The cartoons depict the two experimental strategies. (D) L: Crude lysate; S: supernatant-fraction; P: pellet-fraction. For each experimental condition, the extraction efficiencies (%) of mat-ERj3 or pre-ERj3 are given as percent of the respective intensity as compared to the summed-up intensities of the bands in S plus P.

### Depletion of BiP inhibits import of pre-ERj3 into the ER

The observed failure of Sec63H132Q to rescue the phenotype of Sec63 depletion suggested that ERj3 import into the ER does not only involve Sec63, but also BiP (Fig. 5F, G). SubAB treatment is the method of choice for BiP-depletion in HeLa cells, providing an acute and highly efficient depletion whilst maintaining robust cell viability [19, 20, 23, 38]. Therefore, the effect of BiP-depletion on ER-import of pre-ERj3 was analyzed in SubAB treated HeLa cells under conditions of pre-ERj3 overproduction. An inactive mutant variant of SubAB served as control (SubA_A272_B) [23, 38]. HeLa cells were incubated in the presence of the pre-ERj3 expression plasmid for 25 h. During the last 9 h, SubAB or SubA_A272_B were present and MG132 was absent or present for the last 8 h. Efficient BiP-depletion caused massive accumulation of pre-ERj3 and complete inhibition of mat-ERj3 and ERj3 formation (Fig. 7A). Considering the effects of BiP-depletion (Fig. 7A) and the failed rescue of pre-ERj3 transport by Sec63-H132Q (Fig. 5F, G), the data strongly suggest that BiP and its co-chaperone Sec63 cooperate in the ER-import of pre-ERj3. The comparison of MG132 treated and un-treated samples reiterates that non-transported precursors are degraded by the proteasome.

**Fig. 7.**
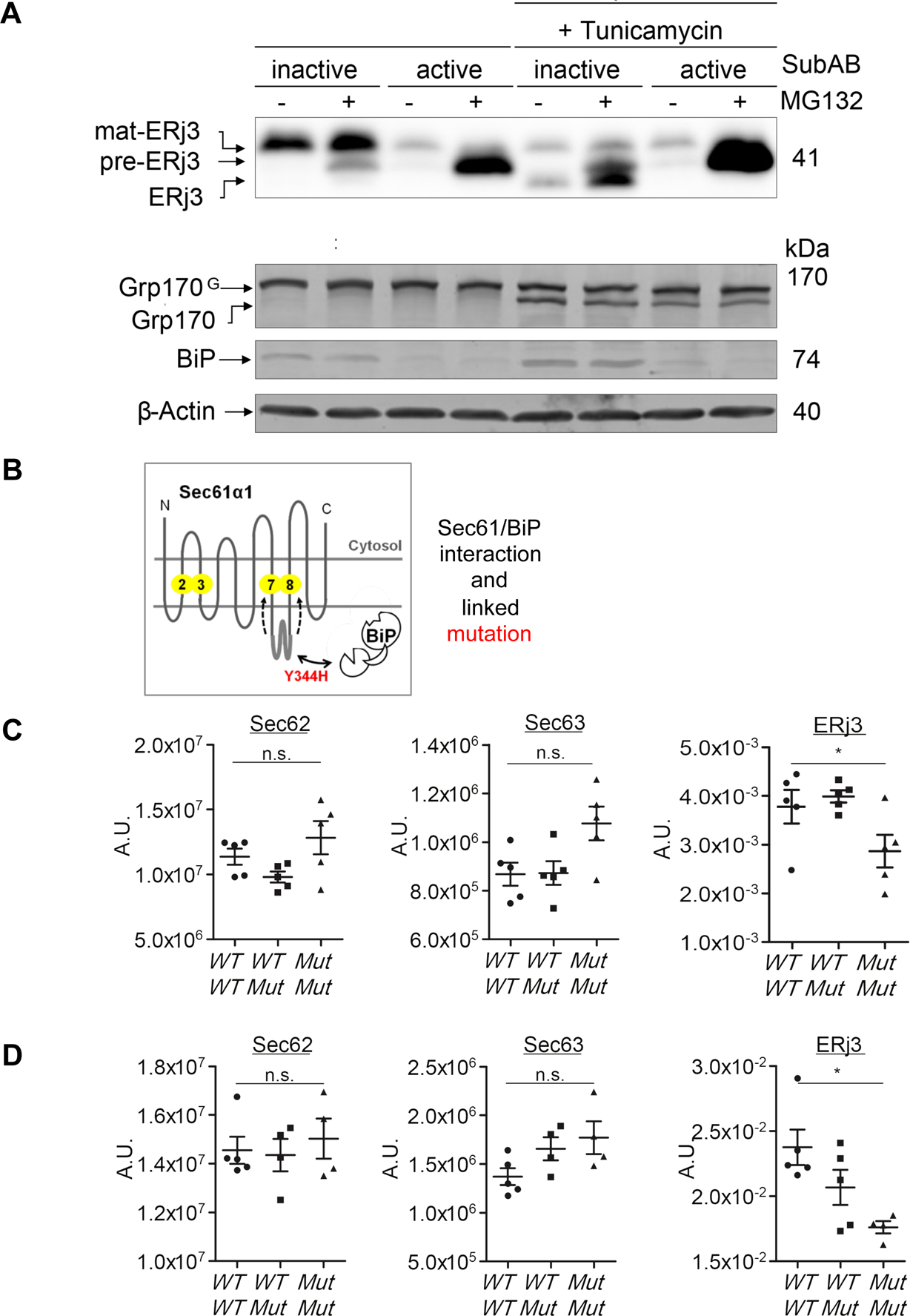
Depletion of BiP from HeLa cells and mutation of the BiP binding site in loop 7 of Sec61α in mice inhibits ER-import of pre-ERj3. (A) Human *ERJ3* was expressed in HeLa cells, which had been treated for 9 h with active or inactive SubAB (1 µg/ml) as indicated, in the presence or absence of MG132 (10 µg/ml) and in the presence or absence of Tunicamycin (2 µg/ml). The consequences of depletion were analyzed by SDS-PAGE and Western blot for the target protein BiP, the model protein ERj3, and the N-glycosylated protein Grp170^G^, ß-actin served as a control. We note that the effect of Tunicamycin can also be deduced from the presence of non-glycosylated Grp170. (B) The cartoon depicts the BiP-interaction with Sec61α1 loop 7 and the Sec61α1 mutant (in red) used in the *in vivo* analysis. (C, D) Dot plots depict relative Sec62-, Sec63-, and ERj3-contents, in liver (C) or pancreas (D) of wildtype mice (WT/WT), heterozygous Sec61α-Y344H mice (WT/Mut), and homozygous Sec61α-Y344H mice (Mut/Mut), respectively, calculated on the basis of SDS-PAGE and Western blots. The mice were of both sexes and various ages. Statistical analysis and graphical representation are described in Experimental Procedures.

A similar set of samples was simultaneously treated with tunicamycin, which inhibits N-glycosylation. BiP-depletion caused accumulation of pre-ERj3 and concomitant inhibition of ERj3 formation, measured as a lack of removal of the SP by ER-lumenal signal peptidase. Therefore, the absence of pre-ERj3 SP removal after BiP-(Fig. 7A), Sec63- and Sec62-depletion (Fig. 5D, F) is consistent with head-on insertion of the SP into the Sec61-channel. In the case of BiP-depletion this assumption was supported by carbonate extraction, too (Fig. 6C, D). Thus pre-ERj3, which accumulated in the absence of BiP may have also been integrated into the ER membrane, consistent with head-on insertion of the SP and a function of all three components in Sec61-channel gating.

Based on *in vitro* ER protein import experiments, the role of BiP in the early stages of import was characterized as a role in Sec61-channel opening, where BiP is recruited to the Sec61-channel via Sec63 and, subsequently, interacts with ER lumenal loop 7 of Sec61α as a substrate (Fig. 7B) [19, 23]. This interaction was suggested to support Sec61-channel gating. Therefore, we addressed the question if this model can be confirmed for pre-ERj3 import into the ER *in vivo*. The BiP binding site in ER lumenal loop 7 of Sec61α is a short helix with a highly conserved twin-tyrosine motif and is sensitive towards the point mutation Y344H, which causes diabetes in homozygous mice with the mutation [19, 23, 39]. Here, we analyzed protein levels of ERj3, Sec62, and Sec63 in pancreas- and liver-tissue from homozygous Sec61α^+/+^, heterozygous Sec61α^+/Y344^ and homozygous Sec61α^Y344H/Y344H^ mice (Fig. 7C, D). The presence of the mutated Sec61α caused reduced ERj3 levels in both organs, while the Sec62 and Sec63 levels were unaffected or even increased. Thus, the BiP-dependence of ERj3 import into the mammalian ER and its action via Sec61α loop 7-interaction were confirmed in different tissues of adult animals.

### The SP plus down-stream positive charges are decisive

To elucidate reasons for the observed dependencies of pre-ERj3 import into the ER of human cells, several mutant variants of pre-ERj3 were tested under the described conditions in HeLa cells. In a first series of experiments, the SP of pre-ERj3 was swapped for a strong SP, the bovine preprolactin (PRL) SP, and, alternatively, two domains of the mature region that are adjacent to the SP were deleted, i.e. either the J-domain (amino acid residues 23-91) or the glycine/phenylalanine-rich-domain (G/F-domain, amino acid residues 92-127) (Fig. 8A) [40]. The three mutant variants were over-produced in comparison to the wild type (WT) precursor in HeLa cells under the various depletion conditions. As expected, depletion of Sec62, Sec63, or BiP caused accumulation of pre-ERj3 and concomitant inhibition of mat-ERj3 formation (Fig. 8B-E). In contrast, replacement of the pre-ERj3 SP by the PRL-SP or deletion of the J-domain, both resulted in reduced dependence of the resulting pre-ERj3 variants on Sec62, Sec63, and BiP. Deletion of the G/F-domain had hardly any effect. However, slightly different phenotypes were observed after depletion of Sec62 and Sec63 as compared to depletion of BiP. While the PRL-SP appeared to almost completely override the Sec62- and Sec63-requirements, it partially retained the requirement for BiP. The deletion of the J-domain, in contrast, almost fully rescued dependencies from the three auxiliary components. Thus, the mature domain, specifically the J-domain following the SP, contributes to the BiP-, Sec63- and Sec62-dependence of pre-ERj3.

**Fig. 8.**
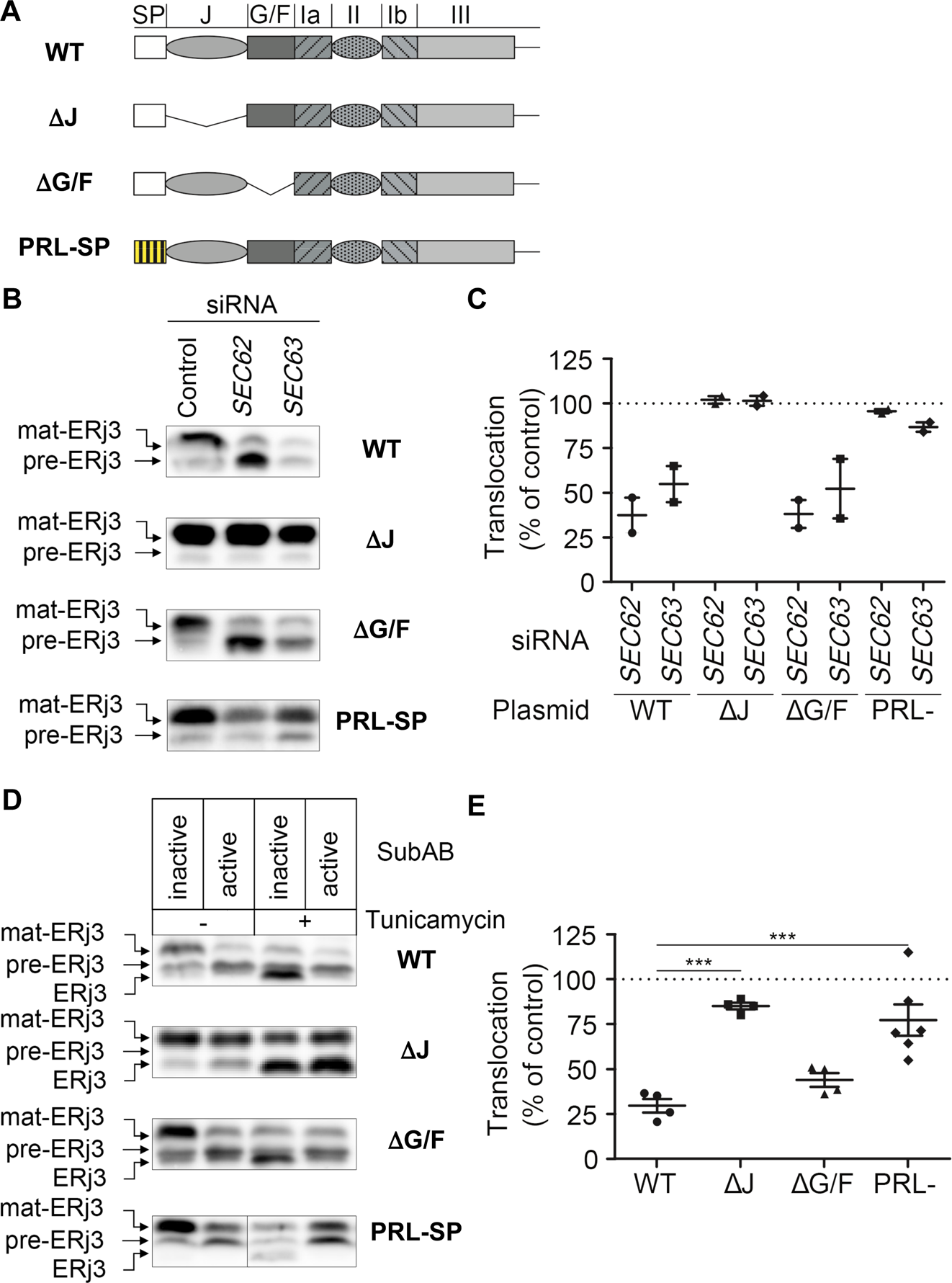
Characterization of pre-ERj3 features as responsible for its dependence on Sec62, Sec63, and BiP - part I. (A-E) Carboxy-terminally HA-tagged murine *ERJ3* or the depicted mutant variants (A) were expressed in HeLa cells, which had been treated with *SEC62*-UTR-targeting siRNA or *SEC63-*UTR-targeting siRNA, or control siRNA (B, C), or active or inactive SubAB (D, E), in the presence of MG132. The consequences of depletion and plasmid complementation were analyzed by SDS-PAGE and Western blot for the model protein ERj3 (anti-HA-tag; anti-ERj3 for PRL SP ERj3-construct). Representative blots are shown in B and D. (C, E) Dot plots depict relative pre-ERj3 translocation efficiencies, calculated as the proportion of N-glycosylation of the total amount of synthesized pre-ERj3 with the individual control sample set to 100%. Statistical analysis and graphical representation are described in Experimental Procedures.

Next, we focussed on the J-domain, which comprises four alpha helices with clusters of positively charged amino acid residues at the amino-termini of helices 2 (_42_KKAYRK) and 4 (_82_KRK) (Fig. 9A). Deletions of helices within the J-domain as well as mutant variants were generated and tested in either CRISPR/Cas9-mediated *SEC62*- and *SEC63*-knock-out HEK293 cells [36] or SubAB treated HeLa cells. In this second series of experiments, either helices 1-3 or 2-3 within the J-domain were deleted or two similar deletions were combined with deletion of the cluster of positively charged amino acid residues at the amino-terminus of helix 4 (Fig. 9A). In addition, the cluster of positively charged amino acid residues at the amino-terminus of helix 2 (_42_KKAYRK) was mutated to either a negative cluster (EEAYEE) or an uncharged one (AAAYAA). These six mutant variants were over-produced in comparison to the wild type (WT) precursor, the SP swap mutant variant, and the J-domain deletion mutant in HEK293 or HeLa cells. More effective depletion of Sec62 or Sec63 in CRISPR/Cas9 treated HEK293 cells caused more effective accumulation of pre-ERj3 in the ER-membrane and concomitant inhibition of ERj3 import into the ER (Fig. 9B, D), as compared to partial depletion (Fig. 8B, C). Depletion of BiP in HeLa cells had the above-described effect (Fig. 9C, E). Under these more effective depletion conditions, replacement of the pre-ERj3 SP by the PRL-SP resulted in partial independence of the resulting pre-ERj3 variant on Sec62, Sec63, and BiP (Fig. 9B-E). Furthermore, deletion of the J-domain resulted in a similarly partial Sec62-independence, but caused an almost complete independence on Sec63 and BiP (Fig. 9B-E). This phenomenon becomes even more pronounced when the other variants are considered. Replacement of the positive cluster within helix 2, _42_KKAYRK, by EEAYEE or AAAYAA did not affect Sec62-dependence (Fig. 9B-E). However, replacement of _42_KKAYRK by AAAYAA caused partial Sec63- and BiP-independence, and replacement of _42_KKAYRK by EEAYEE, i.e. a reversal of charges, caused almost complete Sec63- and BiP-independence. The additional deletion mutant variants were only analyzed for BiP-dependence and led to a striking result. Not only did the two deletion variants of helices 1-3 phenocopy the J-domain deletion, but the movement of the positively charged cluster in helix 4 (_82_KRK) to the previous position of the positively charged cluster of helix 2 (_42_KKAYRK) retained BiP-dependence (∆37-80 variant, Fig. 9C, E). In contrast, this cluster did not cause any harm when it was moved further upstream in the ∆25-80 variant. Thus, it is not only the presence of the positively charged cluster, but the position of this cluster what accounts for BiP-dependence. Furthermore, the inhibitory transport defect of the positive cluster is dominant over the effect of the weak SP and most relevant for Sec63- and BiP-dependence, which represents an almost exact phenocopy of posttranslational ER import of the small presecretory protein preproapelin [19].

**Fig. 9.**
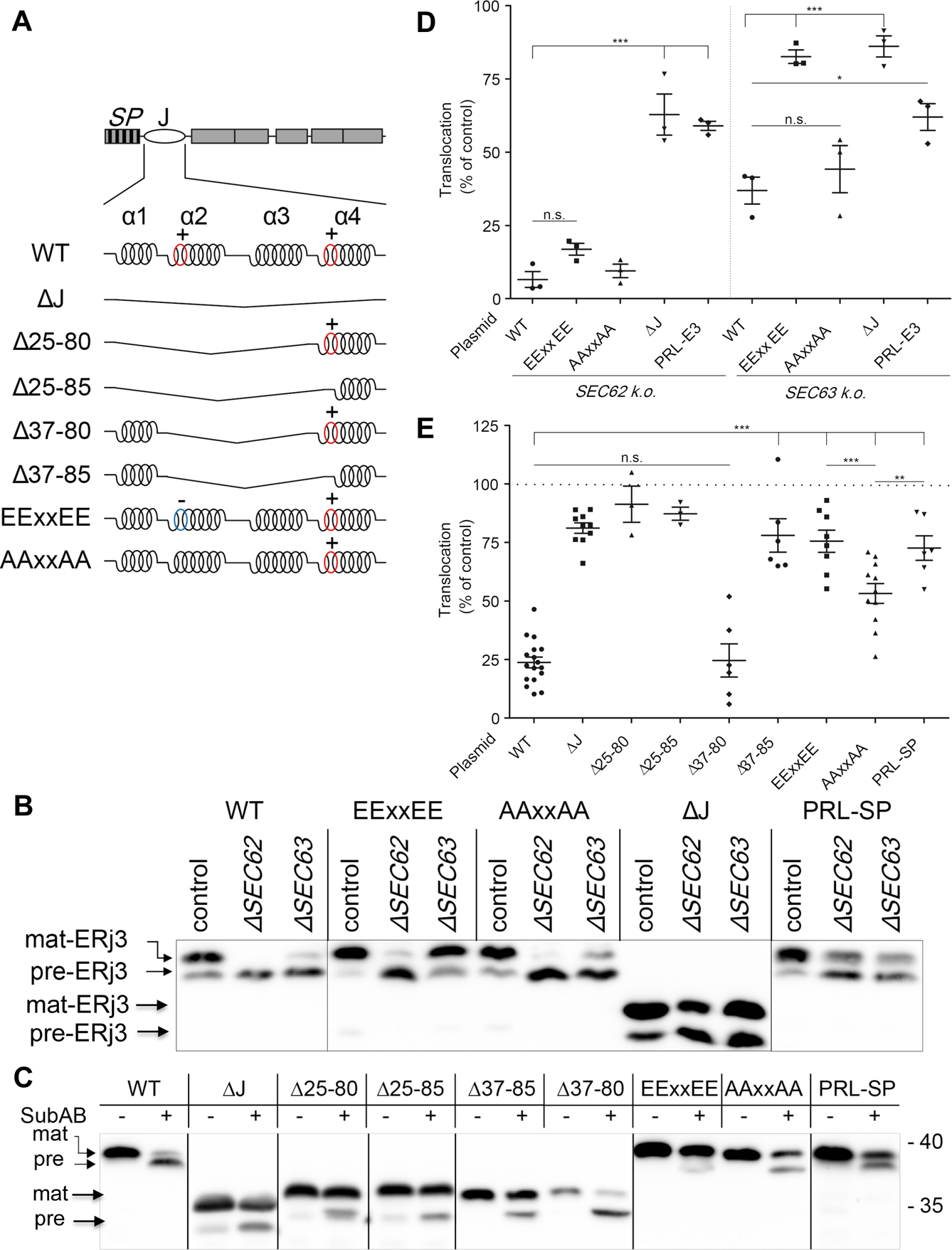
Characterization of pre-ERj3 features as responsible for its dependence on Sec62, Sec63, and BiP – part II. (A-E) Carboxy-terminally FLAG-tagged human *ERJ3* or the depicted mutant variants (**A**) were expressed in HEK293 cells, which had been treated with *SEC62*- or *SEC63*-targeting guide RNA or not treated (B, D), or in HeLa cells, which had been treated with active or inactive SubAB (C, E), all in the presence of MG132. The consequences of depletion and plasmid complementation were analyzed by SDS-PAGE and Western blot for the model protein ERj3 (anti-FLAG-tag). Representative blots are shown in B and C. (D, E) Dot plots depict relative pre-ERj3 translocation efficiencies, calculated as the proportion of N-glycosylation of the total amount of synthesized pre-ERj3 with the individual control sample set to 100%. Statistical analysis and graphical representation are described in Experimental Procedures.

Previous *in vitro* studies of preproapelin suggest that the ability of the CAM741 to inhibit the Sec61-mediated ER-import depends upon the SP plus a positive cluster in the mature region [19, 33]. Furthermore, it was shown that CAM741-sensitivity of preproapelin and its underlying features correlate with its BiP-dependence, which we discussed in a free energy diagram for Sec61-channel gating [19]. We therefore explored the effect of CAM741 on the ER-translocation of pre-ERj3 in intact HeLa cells and found it to be sensitive (Fig. 10A). In contrast, the BiP-independent J-domain deletion variant of pre-ERj3 resulted in a less CAM741-sensitive precursor, providing cellular support for the free energy diagram of Sec61-channel gating (see Discussion).

**Fig. 10.**
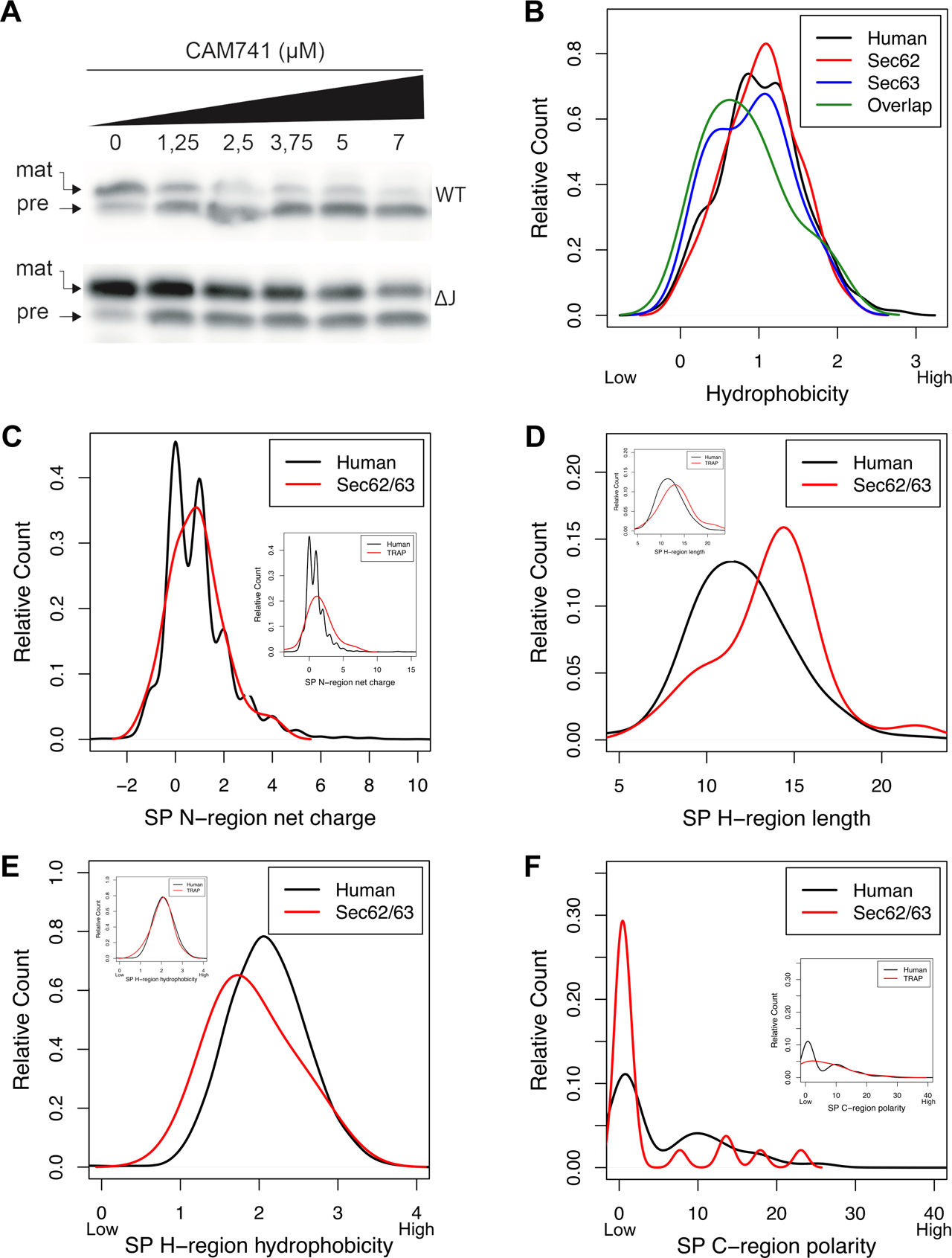
Physicochemical properties of SP and SP-regions of Sec62- and Sec63-clients. (A) Characterization of pre-ERj3 features as responsible for its sensitivity towards the Sec61-channel inhibitor CAM741. HA-tagged murine *ERJ3* or the indicated mutant variant were expressed in HeLa cells, which were treated with DMSO or CAM741 in DMSO at the indicated final concentration for 24 h in the presence of MG132. The consequences of CAM741 treatment were analyzed by SDS-PAGE and Western blot for the model protein ERj3 (anti-HA-tag). (B) Using custom scripts, we computed the hydrophobicity score of SP of Sec62-(n=62) and Sec63-clients (n=21) from HEK293 cells and glycine/proline (GP) content and ∆G_app_ values of the same SP as described in Experimental Procedures. All values were plotted against the relative count. Overlap refers to the SP-comprising clients, which were affected by both manipulations (n=11). Additional plots were computed for TMH. Only hydrophobicity scores of SP showed tendencies and are shown. (C-F) SP segmentation analysis determines the properties of H-, C-, and N-regions of SP of Sec62- and Sec63-clients from HEK293 and HeLa cells (n=23). The regions were determined by segmentation as described in Experimental Procedures. Their displayed properties include N-region net charge (C), H-region length (D), hydrophobicity of H-region (E), and the polarity of C-region (F). Polarity was calculated as the averaged polarity of its amino acids according to the polarity propensity scale. Hydrophobicity was calculated in the same fashion using the Kyte-Doolittle propensity scale. We also used custom scripts to extract all SP annotations for human proteins from UniProtKB entries (Human) and subjected them to the same calculations. The properties of H-, C-, and N-regions of SP of Sec62- and Sec63-clients were also analyzed after their determination by the Phobius (http://phobius.sbc.su.se) prediction tool with similar results. (C-F inserts) For comparison, the SP segmentation analysis was carried out for TRAP clients (21). The values are given in Table S20.

### Characteristics of SP of Sec62/Sec63-dependent precursors

To address the substrate spectrum of the Sec62/Sec63-complex in more detail, we analyzed the data for precursor polypeptides, which were negatively affected by depletion of Sec62 and/or Sec63 in both cell types. Including proteins, which did not meet the significance threshold, the overlap of negatively affected precursor polypeptides between Sec62 and Sec63 depleted HEK293 cells included 27 proteins, 16 with SP and 11 with TMH. Nine of these proteins were below the level of detection in HeLa cells according to this analysis and the literature and, therefore, could not be expected in the overlap of clients between HEK293 and HeLa (Table 2). Taking into account the overlap between *SEC62* and *SEC63* silencing in HeLa cells (ERj3, MAGT1, TGFBI) plus the Sec62 substrates (EPDR1, FKBP10, LNPEP, P4HTM, SLC39A10, TAPBP), a total of 36 precursor polypeptides appeared to be dependent on the Sec62/Sec63-complex in human cells, 23 with SP and 13 with TMH (Table 2). We first analyzed the Sec62- and Sec63-substrates from the HEK293 cells with respect to the physico-chemical properties of their SP and TMH. SP of the 16 Sec62/Sec63-clients showed weak tendencies towards lower than average overall hydrophobicities (Fig. 10B) (Wilcoxon rank test *p*=0.23). However, when next analyzing the N-, H-, and C-regions of all 23 SP (Table 2) a longer than average H-region (*p*=0.01), lower than average H-region hydrophobicity (*p*=0.06), and a lower than average C-region polarity (*p*=0.02) were identified as the distinguishing features for Sec62-plus Sec63-dependence (Fig. 10C-F). A similar phenomenon was not observed for TRAP-dependent precursors [21] (Fig. 10D-F inserts).

## Discussion

Partial or complete translocation through the Sec61-channel in the ER membrane is a crucial step during the biogenesis of about one third of all proteins in eukaryotic cells [1–3]. The opening of this polypeptide-conducting channel during early steps of translocation is mediated by SP and TMH [4–7, 26]. For productive SP or TMH insertion into the Sec61-channel and concomitant opening of the Sec61-channel, a high hydrophobicity/low ∆G^pred^ value for the H-region are conducive [26]. H-region hydrophobicity is recognized by the hydrophobic patch in the Sec61α transmembrane helices 2, 5 and 7, which line the lateral gate of the channel [28]. The SP- and TMH-orientation in the Sec61-channel follows the positive inside rule, i.e. positively charged residues in the N-region support loop insertion, while positively charged side chains downstream of the SP or TMH interfere with loop insertion and favour head-on insertion [26]. For precursor proteins with slowly-gating SP, opening of the mammalian Sec61-channel is supported by allosteric Sec61-channel effectors, i.e. TRAP or the Sec62/Sec63-complex [14–21].

### Sec62 and Sec63 facilitate Sec61-channel opening

This study was aimed at characterizing the function of and rules of engagement for the Sec61-channel effectors Sec62 and Sec63 in human cells. An established unbiased proteomics approach [21] identified a total of 36 precursors as Sec62/Sec63-clients, 23 with SP and 13 with TMH, and, thus, demonstrated at the cellular level that the complex is not only important for posttranslational transport [41]. This may be related to Sec62’s ribosome binding site [37]. The SP-comprising precursors included N-glycoproteins as well as non-glycoproteins and membrane proteins as well as soluble proteins (Table 2). In comparison to all human SP, SP of Sec62/Sec63-clients showed weak tendencies towards lower than average overall hydrophobicities and have longer but less hydrophobic H-regions plus lower C-region polarity. Since H-region hydrophobicity is recognized by the hydrophobic patch in Sec61α [28], we suggest that sampling of SP of the precursors with longer but less hydrophobic H-regions on the ER membrane’s cytosolic face triggers recruitment of the Sec62/Sec63-complex to the Sec61-channel, thereby supporting channel opening [29]. Notably, lower SP hydrophobicity has previously been found to be crucial for Sec62/Sec63-involment in yeast [42]. Therefore, we also subjected SP from *S. cerevisiae* and *E. coli* to segmentation and observed that, in contrast to SP of bacterial precursors, those in yeast are more heterogenous in H-region hydrophobicity, more than half showing a lower hydrophobicity (a value of 2 in average) and the remaining being similar to the majority of human SP (Fig. 11A-D). Strikingly, the SP of the verified SRP-independent and, therefore, Sec72/71/63/62-dependent yeast precursors [42] all fall into the group with lower H-region hydrophobicity (Fig. 11C), just as we have observed here for Sec62/Sec63-clients in human cells (Fig. 10E). Comparison of the shape of the H-region hydrophobicity curves for yeast and human SP suggests that the proportion of Sec62 and Sec63 substrates in human cells is much lower as compared to yeast, i.e. only for yeast the peak shows two distinct maxima, the one with higher H-region hydrophobicity overlapping with the single maximum of human SP. Overall, this is consistent with the comparatively low number of Sec62/Sec63 substrates, which we identified here (Table 1).

**Fig. 11.**
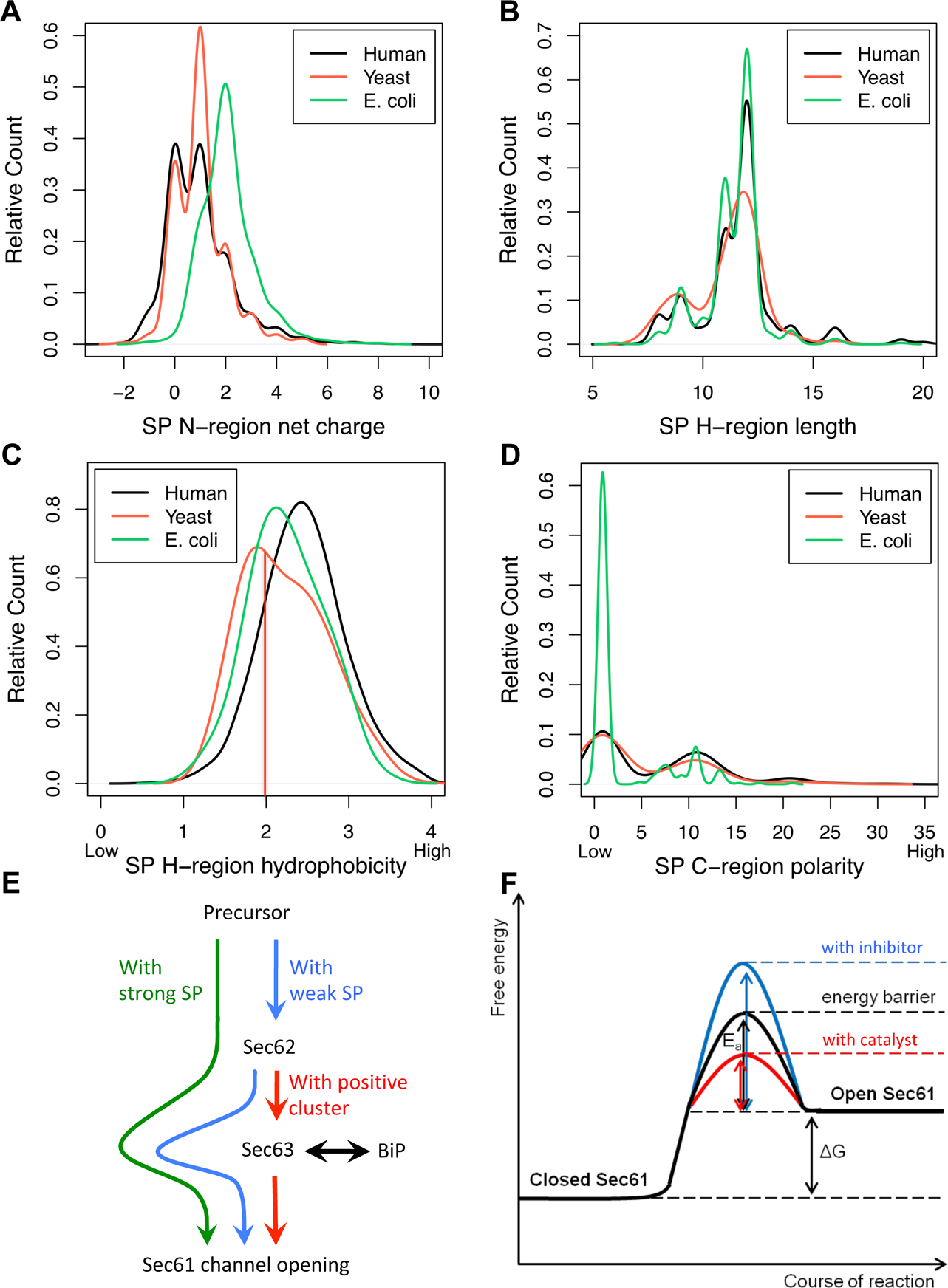
Comparison of physicochemical properties of SP-regions in other species. (A-D) We used custom scripts to extract all SP annotations for human, yeast and *E. coli* proteins from UniProtKB entries. The properties of H-, C-, and N-regions of these SP were analyzed after their determination by the Phobius (http://phobius.sbc.su.se) prediction tool. Their displayed properties include N-region net charge (A), H-region length (B), hydrophobicity of H-region (C), and the polarity of C-region (D). Polarity was calculated as the averaged polarity of its amino acids according to the polarity propensity scale. Hydrophobicity was calculated in the same fashion using the Kyte-Doolittle propensity scale. The values are given in Table S21-23. (E) Requirements for ER-import of pre-ERj3 and their decisive features. (F) Energy diagram for Sec61-channel gating. BiP overcomes the CAM741-reinforced energy barrier for channel opening. E_a_, activation energy; G, free energy.

For one human Sec62/Sec62-client, ERj3, we found that replacing its SP with the strong SP from bovine preprolactin partially overcomes the Sec62/Sec63-dependence, consistent with the SP playing a major but not exclusive role (Fig. 8). As it turned out, a patch of positive charges downstream of the SP, which may force the precursor to follow the positive inside rule in the absence of Sec62/Sec63, contributes to the Sec63-plus the additional BiP-requirement and to a lower extent to the Sec62-requirement (Fig. 11E). However, the additional contribution of the mature region plus the additional BiP-requirement may not be relevant to all Sec62/Sec63-clients and not all of them have such positive patches downstream of the SP. Such patches can e.g. be found in PDIA5 (_19_KKLLRT), RNaseT2 (_2_KRLR), and TGFBI (_15_RLRGR), but not in FKBP10, SDF2, and TMED5 (Fig. 12).

**Fig. 12.**
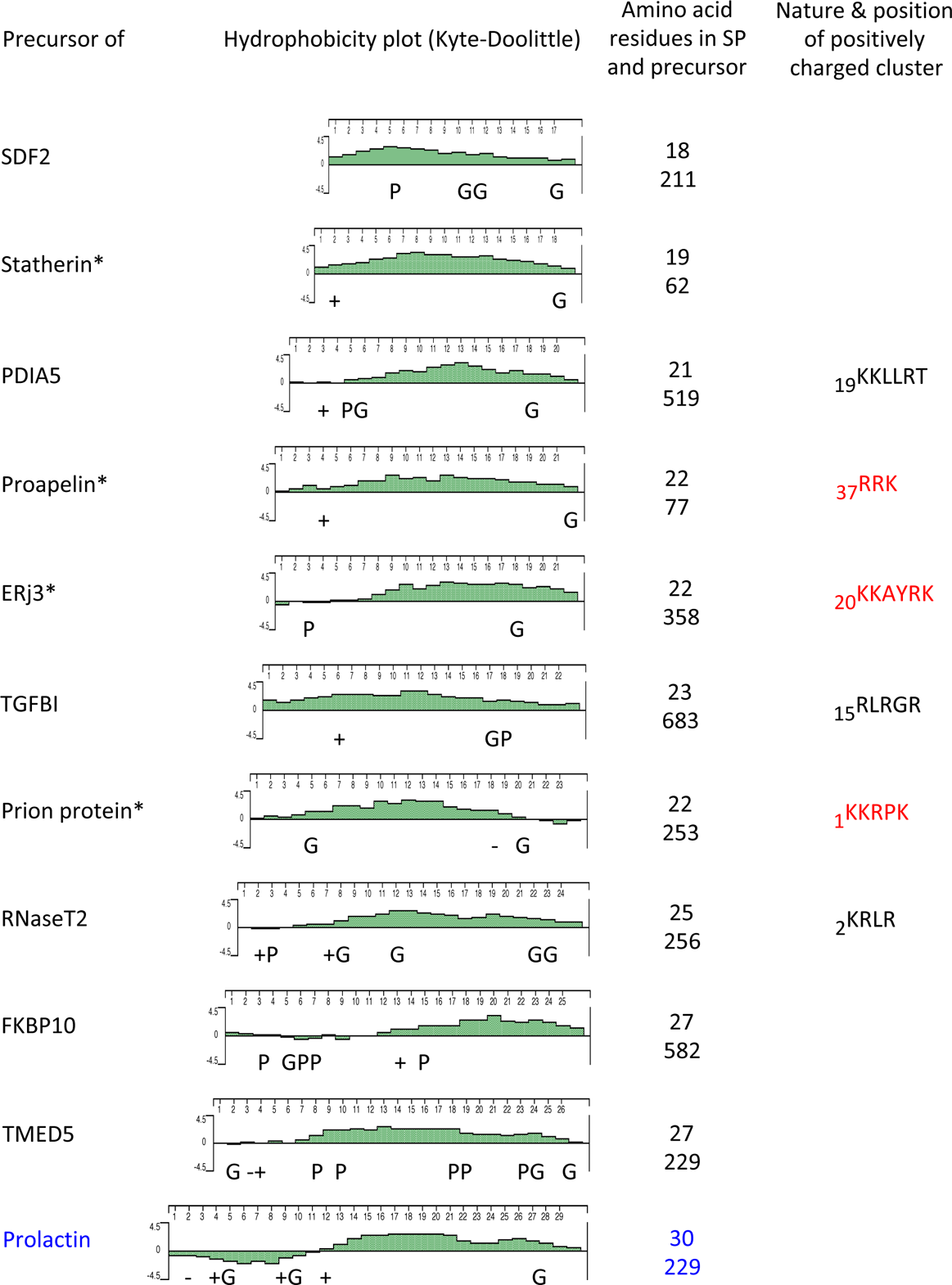
Physicochemical properties of SP and mature parts of selected Sec62- and Sec63- clients. The hydrophobicity analysis of selected SP was carried out using the Protean option of the DNASTAR software package (Lasergene 12). Charged residues (-/+) as well as glycines (G) and prolines (P) are indicated. The total number of amino acid residues of the respective precursor plus its SP is shown. Furthermore, nature and position of positive clusters within the mature domains are given; the clusters, which are shown in red, have experimentally been shown to be decisive. The asterisk indicates precursor polypeptides, which were shown to depend on Sec62 and Sec63 in *in vitro* import assays [19, 20]. The bovine preprolactin SP, shown in blue, is included for comparison.

For the BiP-independent mechanism, the structural analysis of the yeast Sec62/Sec63-complex [24, 25] suggests that Sec63 supports Sec61-channel opening by various interactions, e.g. by wedging of the Sec63 N-terminus between loop 5 of Sec61α and Sec61γ. For the BiP-dependent mechanism, binding of BiP to loop 7 of Sec61α supports Sec61-channel gating to the open state, which may be best interpreted with a free energy diagram for Sec61-channel gating (Fig. 11F) [19, 23]. Both mechanisms are reminiscent of the putative TRAP mechanism, where Sec61-channel opening is mediated by direct interaction of the ER-lumenal domains of the TRAP α- and β-subunits with loop 5 of Sec61α [21]. Low SP hydrophobicity plus long H-region may be the shared features between the respective precursors. Interestingly, nine of the Sec62- or Sec62/Sec63-dependent precursor polypeptides were previously found among the TRAP-dependent proteins after TRAP-depletion [21], possibly hinting at some overlapping but non-identical functions of the two ER membrane protein complexes in Sec61-channel opening (Table 2). This view is consistent with overexpression of the *SSR2* gene under conditions of Sec62-depletion (Fig. 1).

We suggest that non-translocated ERj3 precursor polypeptides accumulating in the absence of Sec62, Sec63, or BiP may incorrectly insert into the ER membrane head-on as an integral membrane protein with type Ia topology (N_ER-lumen_-C_cytosol_). The same phenomenon has previously been reported for the Sec62-client [17] preproinsulin with a SP-mutation (R6C) [43] and -based on *in vitro* experiments-suggested for the posttranslational ER-import of preproapelin [19] and the cotranslational ER-import of the prion protein precursor (Fig. 12) [20]. Strikingly, in the case of the small presecretory protein prestatherin, which lacks a positively charged cluster in the mature region, the Sec62/Sec63-complex was sufficient for efficient translocation [19].

Overall, these results raise the possibility that SP with extended but less hydrophobic H-region depend on a Sec62/Sec63-regulated access to the ER-lumen in human cells. Notably, Sec63 and Sec62 are subject to phosphorylation and Ca^2+^-binding, respectively [44, 45], representing yet another analogy to the TRAP complex [21]. Thus, these modifications are candidates for Sec62/Sec63- and ER protein import-regulation, i.e. the different requirements of different precursors may provide a basis for dual intracellular location of proteins, such as ERj6 (DNAJC3) [46–48], a Sec62-client in HEK293 cells and in HeLa cells, where, however, it did not meet the stringent significance threshold (Table S11).

### Sec61-channelopathies

In the course of the last ten years several human disease were linked to subunits or auxiliary components of the Sec61 complex [5]. These diseases have in common that they affect the gating of the Sec61 channel, i.e. its opening or closing or both, and, therefore, were termed Sec61-channelopathies [49]. They include Common Variable Immune Deficiency (CVID) and Tubolo-Interstitial Kidney Disease (TKD), which were linked to different mutations in the *SEC61A1* gene [50, 51], and Autosomal Dominant PLD and Congenital Disorders of Glycosylation (CDG), which were linked to mutations in the *SEC63* and the *SSR3* and *4* genes, respectively [21, 22, 35]. The heterozygous *SEC61A1-V85D* mutation e.g. decreases ER protein import and increases passive ER calcium efflux via the Sec61-channel, the latter contributing to the short half-live of plasma cells and, therefore, hypogammaglobulinemia [50]. Interestingly, the homozygous *SEC61A1-Y344H* mutation has the same effect on cellular calcium homeostasis but decreases ER protein import in a substrate specific manner and causes the short half-live of pancreatic ß-cells and diabetes in mice [23, 39]. Since Sec63 and the TRAP complex, coded for by the four *SSR* genes, both support Sec61-channel opening in a substrate specific manner, the respective disease linked mutations impair ER import of only a subset of precursors proteins, including certain serum glycoproteins in the case of the TRAP complex and the two plasma membrane proteins polycystin I and II in the case of Sec63 [35]. The reduced level of these two polycystins was proposed to result in the loss of planar cell polarity of cholangiocytes and, thus, cyst formation in the liver. Originally, the latter was attributed to the possible direct role of Sec63 in integration of the two polycystins into the ER membrane. In the light of the results, which were presented here, it is at least as likely that loss of Sec63 function in patient liver cells impairs ER import of ERj3, which represents one of the central players in protein folding in the ER, and that the reduced levels of ERj3 indirectly cause polycystin mis-folding and the disease phenotype. This would easily explain why loss of function mutations in the *PRKCSH* gene, which codes for the ß-subunit of glucosidase II - i.e. a central player in glycoprotein folding in the ER, also cause PLD [35, 52].

## Materials and methods

### Materials

SuperSignal West Pico Chemiluminescence Susbtrate (# 34078) was purchased from Pierce^TM^, Thermo Fisher Scientific. ECL^TM^ Plex goat anti-rabbit IgG-Cy5 (PA45011, used dilution 1:1,000), and ECL^TM^ Plex goat anti-mouse IgG-Cy3 conjugate (PA43009, used dilution 1:2,500) were purchased from GE Healthcare. Horseradish peroxidase coupled anti-rabbit IgG from goat (A 9044, used dilution 1:10,000) and horseradish peroxidase coupled anti-mouse IgG from goat (A 8275, used dilution 1:1,000) were from Sigma-Aldrich. We purchased murine monoclonal antibodies against β-actin (Sigma, A5441, used dilution 1:10,000), HA-tag (Biolegend, 901515, used dilution 1:1,000), the *myc*DDK-tag (Origene, TA50011), and rabbit polyclonal antibodies against FKBP10 (Proteintech, 12172, used dilution 1:1,000), PDIA5 (Proteintech, 15545, used dilution 1:300), RNaseT2 (Proteintech, 13753, used dilution 1:1,000). Additional rabbit antibodies were raised against purified canine proteins (Calreticulin, used dilution 1:250; GRP170, used dilution 1:500), recombinant human protein (Sec63∆N380, used dilution 1:500), the carboxy-terminal peptides of human Sec61α (CKEQSEVGSMGALLF, used dilution 1:250), Sec62 (CGETPKSSHEKS, used dilution 1:500) and ERj3 (CGSVQKVYNGLQGY, used dilution 1:250) with an additional amino-terminal cysteine, and the amino-terminal peptide of BiP (EEEDKKEDVGTVC, used dilution 1:500) with an additional carboxy-terminal cysteine. The antibody directed against ERj3 was affinity purified on Sulfolink-immobilized peptide (Thermo Fisher Scientific). Antibody quality was previously documented [53, 54]. We note that the full scans of the blots are shown in Supplementary Information http://dx.doi.or/10.17632/6s5hn73jcv.1. MG 132 and Tunicamycin were obtained from Calbiochem (# 474790, #654380).

### Animals

Wild type (C57BL/6J) and Sec61α^Y344H^ mutant (C57BL/6J-Sec61a1^m1Gek^/J) mice were purchased from The Jackson Laboratory to establish a colony in our SPF animal facility [39]. Mouse genotyping was carried out on ear punches by Eurofins genomics. Animals had free access to tap water, were fed with standard chow (V1185-300 during breeding, otherwise V1534-300, ssniff Spezialitäten GmbH), and sacrificed for organ collection at various ages by an overdose of anesthetics (ketamine [Ursotamin™, Serumwerk Bernburg] plus xylazine [Rompun™, Bayer]). All animal experiments were performed in accordance with the German legislation on protection of animals (§ 8 TierSchG), the EU Directive 2010/63/EU and the National Institutes of Health Guidelines for the Care and Use of Laboratory Animals (NIH Publication #85-23 Rev. 1985) and were approved by the local governmental animal care committee (approval number 25/2014).

### Cell manipulations

HeLa cells (DSM no. ACC 57) were obtained from the German Collection of Microorganisms and Cell Cultures, routinely tested for mycoplasma contamination by VenorGeM Mycoplasm Detection Kit (Biochrom AG, WVGM), and replaced every five years by a new batch. They were cultivated at 37°C in a humidified environment with 5% CO_2_, in DMEM with 10% fetal bovine serum (FBS; Sigma-Aldrich) and 1% penicillin and streptomycin.

For transient gene silencing, 5.2×10^5^ HeLa cells were seeded per 6-cm culture plate, followed by incubation under normal culture conditions [18]. For *SEC62* or *SEC63* silencing, the cells were transfected with a final concentration of 20 nM targeting siRNA (Qiagen) or 20 nM AllStars Negative Control siRNA (Qiagen) using HiPerFect Reagent (Qiagen) following the manufacturer’s instructions. The targeting siRNAs had the following sequences: *SEC62*-siRNA, GGCUGUGGCCAAGUAUCUUdTdT; *SEC62*-UTR-siRNA, CGUAAAGUGUAUUCUGUACdTdT; *SEC63-*UTR*-*siRNA#1, GGGAGGUGUAGUUUUUUUAdTdT; *SEC63-*UTR-siRNA#2, CAGCUUUAGUUUUAGCAAAdTdT). After 24 h, the medium was changed and the cells were transfected a second time. In each case, silencing was performed for a total of 96 h. BiP-depleted cells were obtained by treating HeLa cells with the subtilase cytotoxin SubAB, which specifically inactivates BiP, at a final concentration of 1 µg/ml for 9 h [23, 38]. Control cells were treated with SubA_A272_B, an inactive mutant form of SubAB.

To rescue the phenotype after *SEC62* or *SEC63* silencing the corresponding human cDNAs, were inserted into the multi-cloning sites of a pCDNA3-IRES-GFP [55]. Cells were treated with *SEC62-*UTR or *SEC63*-UTR siRNA as described above for 96 h. Six hours after the second transfection, the siRNA treated cells were transfected with either vector or expression plasmid using Fugene HD (Promega).

For construction of the guideRNA-Cas9 plasmid, lentiCRISPRv2-puro system (Addgene 52961) was obtained from Addgene [36]. The target sequences for guide RNA were synthetized by Microsynth and corresponded to +242 to +261 and +83 to +102 nucleotides from the transcriptional start site of human Sec62 (GI: 1928972) and Sec63 (GI:3978516), respectively. Two annealed oligonucleotides were inserted into lentiCRISPRv2-puro vector using the BsmBI restriction site. The plasmid was transfected with Jet Prime (Polyplus) into Flp-In^TM^ T-REx^TM^ HEK293 inducible cells (Invitrogen) according to the manufacturer’s instructions to generate CRISPR lines. The cells were cultured in DMEM supplemented with 10% FBS, 100 µg/ml zeocin and 15 µg/ml blasticidin. Two days after transfection, the medium was changed with addition of 2 µg/mL puromycin. Puromycin-resistant clones were picked after 10 days.

### Cell analysis

Growth rates and viability were determined by employing the Countess^®^ Automated Cell Counter (Invitrogen). Silencing efficiencies were evaluated by Western blot using the respective antibodies and an anti-β-actin antibody for sample comparison. Primary antibodies were visualized with ECL^TM^ Plex goat anti-rabbit IgG-Cy5 conjugate (for BiP, Grp170, Sec62, Sec63), or ECL^TM^ Plex goat anti-mouse IgG-Cy3 conjugate (ß-actin) using the Typhoon-Trio imaging system combined with Image Quant TL software 7.0 (GE Healthcare). Alternatively, peroxidase coupled anti-rabbit IgG (for BiP, Calreticulin, ERj3, FKBP10, Grp170, PDIA5, RNaseT2, Sec61α, Sec62, Sec63), or peroxidase coupled anti-mouse IgG (for FLAG-tag, HA-tag) were employed in combination with SuperSignal West Pico Chemiluminescent Substrate and the Fusion SL (peqlab) luminescence imaging system with accompanying software.

For plasmid driven overproduction of model precursor polypeptide pre-ERj3 and its mutant variants, HeLa cells were cultured in the presence of siRNA for a total of 96 h. After 72 h, the cells were transfected with either i) the pCDNA3-IRES-GFP-vector, comprising the human *ERJ3* cDNA [55] or its mutant variant (PRL SP ERj3-construct), or ii) the pCMV6AC-IRES-GFP-vector, comprising the human *ERJ3* cDNA or its mutant variants with an additional carboxy-terminal FLAG-tag (∆J, PRL SP ERj3, and all site directed mutants), or iii) the HA-DSL vector, comprising the murine *ERJ3* cDNA or its deletion mutant variants with an additional carboxy-terminal HA-tag (∆J and ∆G/F) [40], using Fugene HD. After 88 h, Tunicamycin (2 µg/ml) and/or MG 132 (10 µM) were added where indicated. The human mutant variants were generated by SOE-PCR and cloned into the pCMV6AC-IRES-GFP-vector. All constructs were validated by DNA sequencing. Notably, human (GenBank: CAG33377.1) and murine (GenBank: AAH40747.1) pre-ERj3 are 98% identical at the amino acid sequence level.

Semi-permeabilized cells were prepared from equal cell numbers by washing in PBS, and subsequent treatment with digitonin for 5 min at 0 °C^18^. A pellet of 20 × 10^6^ cells was resuspended in 1 ml ice-cold KHM buffer (110 mM potassium acetate, 2 mM magnesium acetate in 20 mM HEPES/KOH, pH 7.2) and supplemented with 22 µl digitonin stock solution (40 mg/ml DMSO). Where indicated, sequestration assays were performed with 50 µl aliquots for 60 min at 0°C in 80 mM sucrose supplemented or not with combinations of proteinase K (50 µg/ml) plus trypsin (50 µg/ml) and Triton-X100 (0.1%) or H_2_O as indicated. Proteolysis was stopped by the addition of phenylmethylsulphonyl fluoride (final concentration: 20 mM) and incubation continued for 5 min at 0°C. Alternatively, semi-permeabilized cells were subjected to alkaline extraction. For this, cells from 100 µl aliquots were re-isolated by centrifugation, resuspended in 100 mM sodium carbonate (pH 11.5), and incubated for 1 h at 4 °C. Subsequently, the solution was subjected to centrifugation for 1 h at 200,000 x g and 2 °C to separate the extracted from integral membrane proteins.

### Label-free quantitative proteomic analysis

After growth for 96 h, 1 × 10^6^ cells (corresponding to roughly 0.2 mg protein) were harvested, washed twice in PBS, and lysed in buffer containing 6 M GnHCl, 20 mM tris(2-carboxyethyl)phosphine (TCEP; Pierce^TM^, Thermo Fisher Scientific), 40 mM 2-chloroacetamide (CAA; Sigma-Aldrich) in 100 mM Tris, pH 8.0 [21]. The lysate was heated to 95°C for 2 min, and then sonicated in a Bioruptor sonicator (Diagenode) at the maximum power setting for 10 cycles of 30 s each. For a 10% aliquot of the sample, the entire process of heating and sonication was repeated once, and then the sample was diluted 10-fold with digestion buffer (25 mM Tris, pH 8, 10% acetonitrile). Protein extracts were digested for 4 h with Lysyl endoproteinase Lys-C (Wako Bioproducts, enzyme to protein ratio: 1:50), followed by the addition of trypsin (Promega) for overnight digestion (enzyme to protein ratio: 1:100). The next day, booster digestion was performed for 4 h using an additional dose of trypsin (enzyme to protein ratio: 1:100). After digestion, a 10% aliquot of peptides (corresponding to about 2 µg of peptides) were purified via SDB-RPS StageTips [56], eluted as one fraction, and loaded for mass spectrometry analysis. Purified samples were loaded onto a 50-cm column (inner diameter: 75 microns; packed in-house with ReproSil-Pur C18-AQ 1.9-micron beads, Dr. Maisch GmbH) via the autosampler of the Thermo Easy-nLC 1000 (Thermo Fisher Scientific) at 60°C. Using the nanoelectrospray interface, eluting peptides were directly sprayed onto the benchtop Orbitrap mass spectrometer Q Exactive HF (Thermo Fisher Scientific) [57]. Peptides were loaded in buffer A (0.1% (v/v) formic acid) at 250 nl/min and the percentage of buffer B was ramped to 30% over 180 min, followed by a ramp to 60 % over 20 min, then 95 % over the next 10 min, and maintained at 95% for another 5 min [58]. The mass spectrometer was operated in a data-dependent mode with survey scans from 300 to 1,700 m/z (resolution of 60,000 at m/z = 200). Up to 15 of the top precursors were selected and fragmented using higher energy collisional dissociation (HCD) with a normalized collision energy value of 28 [59]. The MS2 spectra were recorded at a resolution of 17,500 (at m/z = 200). AGC target for MS and MS2 scans were set to 3E6 and 1E5, respectively, within a maximum injection time of 100 and 25 ms for MS and MS2 scans, respectively. Dynamic exclusion was enabled to minimize repeated sequencing of the same precursor ions and set to 30 s [59].

Raw data were processed using the MaxQuant computational platform [60]. The peak list was searched against Human Uniprot databases, with an initial precursor mass deviation up to 4.5 ppm for main search and an allowed fragment mass deviation of 20 ppm [58]. MaxQuant by default enables individual peptide mass tolerance and was used in the search. Cysteine carbamidomethylation was set as the static modification, and methionine oxidation and N-terminal acetylation as variable modifications. The match between the run feature was enabled, and proteins were quantified across samples using the label-free quantification algorithm in MaxQuant [61] as label-free quantification (LFQ) intensities. Notably, LFQ intensities do not reflect true copy numbers because they depend not only on the amounts of the peptides but also on their ionization efficiencies; thus, they only served to compare abundances of the same protein in different samples [60–62]. The mass spectrometry proteomics data have been deposited to the ProteomeXchange Consortium via the PRIDE [63] partner repository at http://www.ebi.ac.uk/pride/archive/projects/Identifiers with the dataset identifiers: PXD008178, PXD011993, and PXD012078.

### Experimental Design and Statistical Rational for Proteome Data

Data processing was carried out as previously described [21]. Each HeLa cell MS experiment contained three groups of samples: one control and two silencing experiments (down-regulation by two different siRNAs). Each group consisted of three data points (replicates). All proteins having only one or zero valid data points for the control condition were neglected. For Sec62, only proteins that were detected in both silencing experiments were considered. Missing data points were generated by imputation [21], whereby we distinguished two cases. For proteins that were completely missing (lacking any valid data point) in one silencing condition, imputed data points were randomly generated in the bottom tail of the whole proteomics distribution following the strategy of the Perseus software (http://www.biochem.mpg.de/5111810/perseus) [64]. For proteins having at least one valid MS data point in one condition, missing data points were generated with the local least squares (LLS) imputation method [65]. Gene-based quantile normalization was applied to homogenize the abundance distributions of each protein with respect to statistical properties. To identify which proteins were affected by the knock-down of the targeted proteins (Sec62 and Sec63) in siRNA-induced cells, we log2-transformed the ratio between siRNA and control samples and conducted 2 separate t-tests for each siRNA against the control sample [21]. The p-value was corrected by a permutation false discovery rate (FDR) test. Proteins with FDR lower than 5% were considered significantly affected by the silencing of the targeted proteins. Afterwards, the results from the two t-tests were intersected for further analysis. All statistical analysis was done using the R package SAM (http: www-stat-class.stanford.edu) [66]. The HEK293 cell MS experiments were analyzed similarly.

Protein annotations of SP, transmembrane regions, and N-glycosylation sites in humans and yeast were extracted from UniProtKB entries using custom scripts. Using custom scripts, we computed the hydrophobicity score and glycine/proline (GP) content of SP and TMH sequences. A peptide’s hydrophobicity score was assigned as the average hydrophobicity of its amino acids according to the Kyte-Doolittle propensity scale (averaged over the sequence length) [67]. GP content was calculated as the total fraction of glycine and proline in the respective sequence [21]. ∆G_app_ values of SP and TMH were calculated with the ∆G_app_ predictor for TM helix insertion (http://dgpred.cbr.su.se).

SP segmentation prediction was carried out using two alternative approaches. In the first approach, we used the well-established prediction tool Phobius [68] to identify N-region, H-region and C-region of all SP (n=2876). Based on this, we calculated the total net charge of the N-region, the polarity of the C-region, and the hydrophobicity and absolute length of the H-region. The polarity score of a single peptide was calculated as the averaged polarity of its amino acids according to the polarity propensity scale derived by Zimmerman et al. [69]. The hydrophobicity score was calculated in the same fashion using the Kyte-Doolittle propensity scale [67]. In the second approach (n=3528), we adopted the sliding window method described by Kyte-Doolittle [67] to calculate a window-averaged hydrophobicity score for each amino acid (n=4 residues on each side around the central residue). A stretch of continuous amino acids which all have a window-averaged hydrophobicity score higher than a certain threshold is considered as the H-region of the SP. An optimal hydrophobicity threshold was derived by applying the same sliding window method to the central 50% portion of all transmembrane domains (TMD set) and all signal peptides (SP set) annotated for the human proteome in Uniprot. The threshold 0.41 gave the best agreement of mean and variance between SP set and TMD set. Around the H-region identified for an SP with this optimal threshold, the remaining upstream and downstream portions of the SP were considered as N-region and C-region, respectively. The physicochemical properties of the C-, H- and N-regions were then analyzed similarly as for the first approach.

### Experimental Design and Statistical Rational for other Data

Dot plots depict relative protein amounts or translocation efficiencies, calculated as the proportion of N-glycosylation and/or SP cleavage of the total amount of synthesized precursors with the individual control sample set to 100%. Data points, the mean of at least three independent experiments, and the standard error of the mean were visualized with GraphPad Prism 5 software. For statistical comparison between a treatment group and the corresponding control a Student’s t-test was used. To compare between multiple precursor variants or treatment groups one-way ANOVA was performed including the post hoc Dunnett or Newman-Keuls test, respectively, using normalized values. Significance levels are given as follows: p < 0.001 (***), p < 0.01 (**), p < 0.05 (*).

## Author Contributions

S.S. performed all ERj3 experiments. J.D. and R.Z. planned and supervised the sample generation for MS analysis. N.N. carried out the MS analysis. D.N. performed the MS data analysis and analyzed SP characteristics under supervision by V.H. S.H. performed the validating Western blots. A.C. and P.W. coordinated animal breeding and organ collection. M.G. supervised the animal genotyping. A.W.P. and J.C.P. provided purified wild type and mutant SubAB, M.L. and M.M. the different HEK293 cells. F.F., J.D., and R.Z. designed the study and wrote the manuscript together with S.L. and V.H. All authors discussed the results and the manuscript.

## Acknowledgements

We are thankful to Monika Lerner and Aline Herges (Homburg) for technical assistance in MS sample generation and organ collection, respectively, to Linda Hendershot (St. Jude Children’s Research Hospital, Memphis, USA) for the murine ERj3 coding plasmids, to Sorin V. Fedeles and Stefan Somlo (Yale School of Medicine, New Haven, USA) for the murine *SEC63^-/-^* cells, to Novartis (Basel, Switzerland) for CAM741, and to Martin van der Laan (Homburg) for financial support for S.H..

## Funding sources and disclosure of conflicts of interest

This research was supported by the DFG grants ZI234/13-1, FO716/4-1. M.M. was supported by Signora Alessandra, AlphaONE Foundation, Foundation for Research on Neurodegenerative Diseases, Swiss National Science Foundation (SNF), and Comel and Gelu Foundations. The authors declare that they have no conflict of interest.

## Supporting information

Table S1_sec62_all.xlsx

Complete list of genes corresponding to proteins quantified in both Sec62 depletion experiments. Gene names, protein accession numbers (ID), log2 fold changes resulting from siRNA-mediated Sec62 depletion, and -log10 p values are indicated. Minus sign in front of fold change denotes negatively affected proteins. The number of listed proteins differs from the total number of quantified proteins because some proteins were not quantified in both experiments, or were quantified in less than two of the triplicates in at least one experiment. The original Orbitrap data for all quantified proteins are deposited at Proteome Exchange: http://www.proteomexchange.org.

Table S2_sec62_full_lo.xlsx

Proteins that were negatively affected by Sec62 depletion, i.e. putative Sec62 clients. Gene names, protein accession numbers, and log2 fold changes resulting from siRNA-mediated Sec62 depletion are presented together with full protein names and Gene Ontology (GO) annotations for subcellular location(s), as extracted from UniProtKB entries using custom scripts. Proteins are listed according to decreasing negative effects of Sec62 depletion.

Table S3_sec62_full_up.xlsx

Proteins that were positively affected by Sec62 depletion. Gene names, protein accession numbers, and log2 fold changes resulting from siRNA-mediated Sec62 depletion are presented together with Gene Ontology (GO) annotations for subcellular location(s), as extracted from UniProtKB entries using custom scripts. Proteins are listed according to decreasing positive effects of Sec62 depletion.

Table S4_sec62_full_lo_2.xlsx

Proteins that were negatively affected by siRNA-mediated Sec62 depletion and comprise signal peptides or transmembrane helices, which are serving as signal peptides, i.e. Sec62 clients. Gene names, protein accession numbers, and log2 fold changes resulting from Sec62 depletion are presented together with presence of signal peptide (SP) or transmembrane helix (TMH), total number of transmembrane helices (TM regions), number of N-glycosylation sites (N-glyc sites), amino acid sequences of SP or TMH (in single letter code), TMH position within the total amino acid sequence (all as extracted from UniProtKB entries using custom scripts) and glycine plus proline content of SP or TMH (GP%), hydrophobicity (Hph), predicted delta G (all as described in Methods). Proteins are listed according to decreasing negative effects of Sec62 depletion.

Table S5_segmentation_sec62_full_lo_3_sp.xlsx

Segmentation analysis of SP of Sec62 clients. Gene names are presented together with N-region net charge, N-region length, H-region hydrophobicity, H-region length, C-region polarity after SP segmentation by the Phobius (http://phobius.sbc.su.se) prediction tool, and total glycine plus proline content of SP (per_gp), overall hydrophobicity, predicted delta G (delta_g), and amino acid sequences (in single letter code) (all as described in Methods).

Table S6_sec63_all.xlsx

Complete list of genes corresponding to proteins quantified in the Sec63 depletion experiment. Gene names, protein accession numbers (ID), log2 fold changes resulting from siRNA-mediated Sec63 depletion, and -log10 p values are indicated. Minus sign in front of fold change denotes negatively affected proteins. The number of listed proteins differs from the total number of quantified proteins because some proteins were quantified in less than two of the triplicates. The original Orbitrap data for all quantified proteins are deposited at Proteome Exchange: http://www.proteomexchange.org.

Table S7_sec63_full_lo.xlsx

Proteins that were negatively affected by Sec63 depletion, i.e. putative Sec63 clients. Gene names, protein accession numbers, and log2 fold changes resulting from siRNA-mediated Sec63 depletion are presented together with full protein names and Gene Ontology (GO) annotations for subcellular location(s), as extracted from UniProtKB entries using custom scripts. Proteins are listed according to decreasing negative effects of Sec63 depletion.

Table S8_sec63_full_up.xlsx

Proteins that were positively affected by Sec63 depletion. Gene names, protein accession numbers, and log2 fold changes resulting from siRNA-mediated Sec63 depletion are presented together with Gene Ontology (GO) annotations for subcellular location(s), as extracted from UniProtKB entries using custom scripts. Proteins are listed according to decreasing positive effects of Sec63 depletion.

Table S9_sec63_full_lo_2.xlsx

Proteins that were negatively affected by siRNA-mediated Sec63 depletion and comprise signal peptides or transmembrane helices, which are serving as signal peptides, i.e. Sec63 clients. Gene names, protein accession numbers, and log2 fold changes resulting from Sec63 depletion are presented together with presence of signal peptide (SP) or transmembrane helix (TMH), total number of transmembrane helices (TM regions), number of N-glycosylation sites (N-glyc sites), amino acid sequences of SP or TMH (in single letter code), TMH position within the total amino acid sequence (all as extracted from UniProtKB entries using custom scripts) and glycine plus proline content of SP or TMH (GP%), hydrophobicity (Hph), predicted delta G (all as described in Methods). Proteins are listed according to decreasing negative effects of Sec63 depletion.

Table S10_sec62_crispr_all.xlsx

Complete list of genes corresponding to proteins quantified in the Sec62 depletion experiment. Gene names, protein accession numbers (ID), log2 fold changes resulting from CRISPR/Cas9-mediated Sec62 depletion, and -log10 p values are indicated. Minus sign in front of fold change denotes negatively affected proteins. The number of listed proteins differs from the total number of quantified proteins because some proteins were quantified in less than two of the triplicates. The original Orbitrap data for all quantified proteins are deposited at Proteome Exchange: http://www.proteomexchange.org.

Table S11_sec62_crispr_full_lo.xlsx

Proteins that were negatively affected by Sec62 depletion, i.e. putative Sec62 clients. Gene names, protein accession numbers, and log2 fold changes resulting from CRISPR/Cas9-mediated Sec62 depletion are presented together with full protein names and Gene Ontology (GO) annotations for subcellular location(s), as extracted from UniProtKB entries using custom scripts. Proteins are listed according to decreasing negative effects of Sec62 depletion.

Table S12_sec62_crispr_full_up.xlsx

Proteins that were positively affected by Sec62 depletion. Gene names, protein accession numbers, and log2 fold changes resulting from CRISPR/Cas9-mediated Sec62 depletion are presented together with Gene Ontology (GO) annotations for subcellular location(s), as extracted from UniProtKB entries using custom scripts. Proteins are listed according to decreasing positive effects of Sec62 depletion.

Table S13_sec62_crispr_full_lo_2.xlsx

Proteins that were negatively affected by CRISPR/Cas9-mediated Sec62 depletion and comprise signal peptides or transmembrane helices, which are serving as signal peptides, i.e. Sec62 clients. Gene names, protein accession numbers, and log2 fold changes resulting from Sec62 depletion are presented together with presence of signal peptide (SP) or transmembrane helix (TMH), total number of transmembrane helices (TM regions), number of N-glycosylation sites (N-glyc sites), amino acid sequences of SP or TMH (in single letter code), TMH position within the total amino acid sequence (all as extracted from UniProtKB entries using custom scripts) and glycine plus proline content of SP or TMH (GP%), hydrophobicity (Hph), predicted delta G (all as described in Methods). Proteins are listed according to decreasing negative effects of Sec62 depletion.

Table S14_segmentation_sec62_crispr_full_lo_3_sp.xlsx

Segmentation analysis of SP of Sec62 clients. Gene names are presented together with N-region net charge, N-region length, H-region hydrophobicity, H-region length, C-region polarity after SP segmentation by the Phobius (http://phobius.sbc.su.se) prediction tool, and total glycine plus proline content of SP (per_gp), overall hydrophobicity, predicted delta G (delta_g), and amino acid sequences (in single letter code) (all as described in Methods).

Table S15_sec63_crispr_all.xlsx

Complete list of genes corresponding to proteins quantified in the Sec63 depletion experiment. Gene names, protein accession numbers (ID), log2 fold changes resulting from CRISPR/Cas9-mediated Sec63 depletion, and -log10 p values are indicated. Minus sign in front of fold change denotes negatively affected proteins. The number of listed proteins differs from the total number of quantified proteins because some proteins were quantified in less than two of the triplicates. The original Orbitrap data for all quantified proteins are deposited at Proteome Exchange: http://www.proteomexchange.org.

Table S16_sec63_crispr_full_lo.xlsx

Proteins that were negatively affected by Sec63 depletion, i.e. putative Sec63 clients. Gene names, protein accession numbers, and log2 fold changes resulting from CRISPR/Cas9-mediated Sec63 depletion are presented together with full protein names and Gene Ontology (GO) annotations for subcellular location(s), as extracted from UniProtKB entries using custom scripts. Proteins are listed according to decreasing negative effects of Sec63 depletion.

Table S17_sec63_crispr_full_up.xlsx

Proteins that were positively affected by Sec63 depletion. Gene names, protein accession numbers, and log2 fold changes resulting from CRISPR/Cas9-mediated Sec63 depletion are presented together with Gene Ontology (GO) annotations for subcellular location(s), as extracted from UniProtKB entries using custom scripts. Proteins are listed according to decreasing positive effects of Sec63 depletion.

Table S18_sec63_crispr_full_lo_2.xlsx

Proteins that were negatively affected by CRISPR/Cas9-mediated Sec63 depletion and comprise signal peptides or transmembrane helices, which are serving as signal peptides, i.e. Sec63 clients. Gene names, protein accession numbers, and log2 fold changes resulting from Sec63 depletion are presented together with presence of signal peptide (SP) or transmembrane helix (TMH), total number of transmembrane helices (TM regions), number of N-glycosylation sites (N-glyc sites), amino acid sequences of SP or TMH (in single letter code), TMH position within the total amino acid sequence (all as extracted from UniProtKB entries using custom scripts) and glycine plus proline content of SP or TMH (GP%), hydrophobicity (Hph), predicted delta G (all as described in Methods). Proteins are listed according to decreasing negative effects of Sec63 depletion.

Table S19_segmentation_sec63_crispr_full_lo_3_sp.xlsx

Segmentation analysis of SP of Sec63 clients. Gene names are presented together with N-region net charge, N-region length, H-region hydrophobicity, H-region length, C-region polarity after SP segmentation by the Phobius (http://phobius.sbc.su.se) prediction tool, and total glycine plus proline content of SP (per_gp), overall hydrophobicity, predicted delta G (delta_g), and amino acid sequences (in single letter code) (all as described in Methods).

Table S20_segmentation_ trap_full_lo_sp.xlsx

Segmentation analysis of SP of TRAP clients. Gene names, protein accession numbers, and log2 fold changes resulting from TRAP depletion are presented together with presence of signal peptide (SP) or transmembrane helix (TMH), total number of transmembrane helices (TM regions), number of N-glycosylation sites (N-glyc sites), amino acid sequences of SP or TMH (in single letter code), TMH position within the total amino acid sequence (all as extracted from UniProtKB entries using custom scripts) and glycine plus proline content of SP or TMH (GP%), hydrophobicity (Hph), predicted delta G (all as described in Methods). Proteins are listed according to decreasing negative effects of TRAP depletion. These previously established SP characteristics^21^ are presented together with N-region net charge, N-region charge, N-region length, H-region hydrophobicity, H-region length, C-region polarity, and C-region length after SP segmentation by the Phobius (http://phobius.sbc.su.se) prediction tool.

Table S21_segmentation_ uniprot_human_sp.xlsx

Segmentation analysis of SP containing proteins in humans. Gene names are presented together with N-region net charge, N-region charge, N-region length, H-region hydrophobicity, H-region length, C-region polarity, C-region length after SP segmentation by the Phobius (http://phobius.sbc.su.se) prediction tool, and total glycine plus proline content of SP (GP), overall hydrophobicity, and predicted delta G (delta_g). Sec62 and/or Sec63 clients are indicated as candidates.

Table S22_segmentation_ uniprot_yeast_sp.xlsx

Segmentation analysis of SP containing proteins in yeast. Gene names are presented together with N-region net charge, N-region charge, N-region length, H-region hydrophobicity, H-region length, C-region polarity, C-region length after SP segmentation by the Phobius (http://phobius.sbc.su.se) prediction tool. SRP-independent proteins according to Ast et al., 2013 (41) are indicated and can be expected to be Sec62 and Sec63 clients in *S. cerevisiae*.

Table S23_segmentation_uniprot_ecoli_sp.xlsx

Segmentation analysis of SP containing proteins in *E. coli*. Gene names are presented together with N-region net charge, N-region charge, N-region length, H-region hydrophobicity, H-region length, C-region polarity, C-region length after SP segmentation by the Phobius (http://phobius.sbc.su.se) prediction tool.

## Notes

http://www.ebi.ac.uk/pride/archive/projects/Identifiers

http://dx.doi.or/10.17632/6s5hn73jcv.1

